# Glutathionylation primes soluble GAPDH for late collapse into insoluble aggregates

**DOI:** 10.1101/545921

**Authors:** M. Zaffagnini, C.H. Marchand, M. Malferrari, S. Murail, S. Bonacchi, D. Genovese, M. Montalti, G. Venturoli, G. Falini, M. Baaden, S.D. Lemaire, S. Fermani, P. Trost

## Abstract

Protein aggregation is a complex physiological process, primarily determined by stress-related factors revealing the hidden aggregation propensity of proteins that otherwise are fully soluble. Here we report a mechanism by which glycolytic glyceraldehyde-3-phosphate dehydrogenase of *Arabidopsis thaliana* (AtGAPC1) is primed to form insoluble aggregates by the glutathionylation of its catalytic cysteine (Cys149). Following a lag phase, glutathionylated AtGAPC1 initiates a self-aggregation process resulting in the formation of branched chains of globular particles made of partially misfolded and totally inactive proteins. GSH molecules within AtGAPC1 active sites are suggested to provide the initial destabilizing signal. The following removal of glutathione by the formation of an alternative disulfide bond between Cys149 and Cys153 reinforces the aggregation process. Besides acting as a protective mechanism against overoxidation, S-glutathionylation of AtGAPC1 triggers an unexpected aggregation pathway with completely different and still unexplored physiological implications.

## INTRODUCTION

A single polypeptide chain may adopt a huge number of different conformations among which only one, or a few, are biologically active. Although native conformations are thermodynamically favored, in some proteins they are separated from non-native conformations by small energy barriers^1, 2^. Misfolded proteins typically expose hydrophobic residues that are buried in native conformations and these residues tend to promote protein aggregation in aqueous environments. Aggregation propensity is highly differentiated among different proteins, depending on their amino acid sequence and post-translational modifications, but still, the capability to aggregate can be considered as an intrinsic property of any type of polypeptide^1, 3^.

Protein aggregates may occur in different shapes^1, 4^. In some cases, protein aggregation starts with the formation of small oligomers and ends up with amyloid fibrils characterized by a typical cross beta-sheet architecture. Several human disorders collectively known as amyloidoses are associated with amyloid depositions made of disease-specific proteins associated in fibrils^5^. Different from amyloid fibrils, protein particulates are insoluble oligomers with a globular shape and no cross beta-sheet architecture as they are formed by proteins that are only partially unfolded^4^. Aggregation-prone proteins tend to form particulates at pH close to their isoelectric point, and amyloid fibrils at pH values in which they bear a strong net charge^6^. Both aggregation products are considered states that virtually any protein can be forced to adopt, though the process might require harsh treatments such as heat or extreme pH values^6^.

The risk of protein aggregation *in vivo* is exacerbated by the high concentration of proteins in living cells^3^. Being at high risk of aggregation, proteins that are particularly abundant tend to adopt conformations that result in higher solubility compared to less abundant ones^7^. Moreover, cells of any domain of life possess a large set of molecular chaperones that limit protein aggregation by shielding the exposed hydrophobic patches of misfolded proteins, thereby promoting their refolding^8–10^. However, in spite of the molecular chaperones and the efficiency of the whole machinery that controls proteome homeostasis, hundreds of proteins remain at high risk of aggregation, even under non-pathologic al conditions. *In vivo*, protein aggregates have been reported in different model organisms including *E. coli^11^*, yeast^12, 13^, *C. elegans*^14, 15^, tomato and tobacco cell cultures^16, 17^, typically as a consequence of aging or heat stress. These protein aggregates are not associated with specific diseases while they may be associated with oxidative stress conditions shared by different types of stress^13^. Oxidative post-translational modifications (Cys and Met oxidation, carbonylation, etc.) have been shown to favour protein aggregation, presumably by lowering the energy barriers separating native from misfolded conformations^18–21^. Although in some cases non-amyloid protein aggregation can induce cell death^22, 23^, protein aggregation may also be beneficial to cells as long as sequestration of aggregates in specific cell sites prevents toxicity^24^. Indeed, non-amyloid protein aggregates may be asymmetrically inherited by daughter cells^25^, dissolved by chaperones^12, 26^ or digested by proteases^27^ or autophagy^28, 29^. All these processes limit cell toxicity. In this sense, protein aggregation may be considered as a last line of defence against stress^2^.

Glyceraldehyde-3-phosphate dehydrogenase (GAPDH) is a ubiquitous and abundant glycolytic enzyme, which was found to aggregate in different types of cells and conditions. *In vitro*, animal GAPDH forms aggregates under strongly oxidative conditions^21–23, 30–34^. The essential catalytic cysteine of GAPDH can be oxidized by H_2_O_2_ to generate a sulfenic acid group that may either react with a second H_2_O_2_ molecule to form a sulfinic acid. Alternatively, the sulfenic acid of the catalytic cysteine can react with a second thiol, like that of reduced glutathione (GSH), to form a mixed disulfide (S-glutathionylation)^35, 36^. While the sulfinic acid cannot be reduced by cell reductants, the glutathionylated cysteine can be reduced back to the thiol group by glutaredoxins or thioredoxins^37, 38^.

The sensitivity of GAPDH to reactive oxygen species (ROS) has important consequences. Since the interaction between cytoplasmic GAPDH (GAPC) and autophagy-related protein 3 (ATG3) negatively regulates autophagy, ROS may induce autophagy in plants by impairing GAPC-ATG3 complex formation^39^. On the other hand, oxidized GAPC activates phospholipase D at the plasma membrane creating a connection between ROS- and lipid-signaling that controls Arabidopsis response to stress^40^. In animal cells, GAPDH sensitivity to H_2_O_2_ was proposed to have a positive role in oxidative stress conditions because it allows rerouting of the primary metabolism from glycolysis to the oxidative pentose phosphate pathway, the resulting NADPH being essential for the antioxidant response, which in turn allows GAPDH recovery^41^. Thanks to its extreme redox sensitivity, both in animals and in plants^42, 43^, GAPDH is now regarded as being a hub of controlled redox responses for metabolic regulation^44^.

Clearly, all functional interactions, catalytic activity and regulatory functions of GAPDH are impaired by aggregation. In Arabidopsis plants, GAPC is suggested to form aggregates in leaves infiltrated with flg22, a pathogen-associated molecular pattern that triggers basal immunity with ROS being implicated in the response^32^. Here we show that Arabidopsis GAPC1 specifically aggregates *in vitro* following oxidation by H_2_O_2_ in the presence of GSH at nearly physiological concentrations. These conditions lead to specific glutathionylation of GAPC1 catalytic cysteines with no other amino acids being modified. Though protected from irreversible oxidation, glutathionylated GAPC1 is conformationally destabilized and, surprisingly, slowly induced to aggregate into oligomeric particles. In the next phase, glutathionylated GAPC1 spontaneously releases GSH and the small oligomeric particles melt into large micrometric clusters made of smaller (pseudo) globular units. Aggregated GAPC1 is partially unfolded and bears a novel disulfide bond engaging the catalytic Cys149 and the conserved Cys153 of the same subunits. Formation of the disulfide bond speeds up the aggregation process. The crystal structure of glutathionylated AtGAPC1 and Molecular Dynamics (MD) calculations derived thereof, provide clues on the mechanism by which a physiological post-translational modification like S-glutathionylation may trigger the collapse of a soluble tetrameric protein into insoluble aggregates.

## RESULTS

### 1. AtGAPC1 may form non-amyloid aggregates

Visible light scattering (turbidity) can be taken as a proxy of protein stability in solution, and turbidity measurements show that native AtGAPC1 in solution (0.2 mg / ml) is stable for hours at room temperature (**Fig. 1a**). Similar behavior was shown in the presence of H_2_O_2_ (125 μM) at 25:1 ratio with AtGAPC1 subunits (**Fig. 1a**), whereas addition of GSH together with H_2_O_2_ (5:1 ratio) caused a dramatic increase in turbidity over time, indicative of protein aggregation (**Fig. 1a**). The increase in turbidity followed a lag-phase of 15-20 min (**Fig. 1a, inset**), and proceeded linearly for more than one hour reaching a plateau in about 2 hours (**Fig. 1a**).

**Figure 1.**
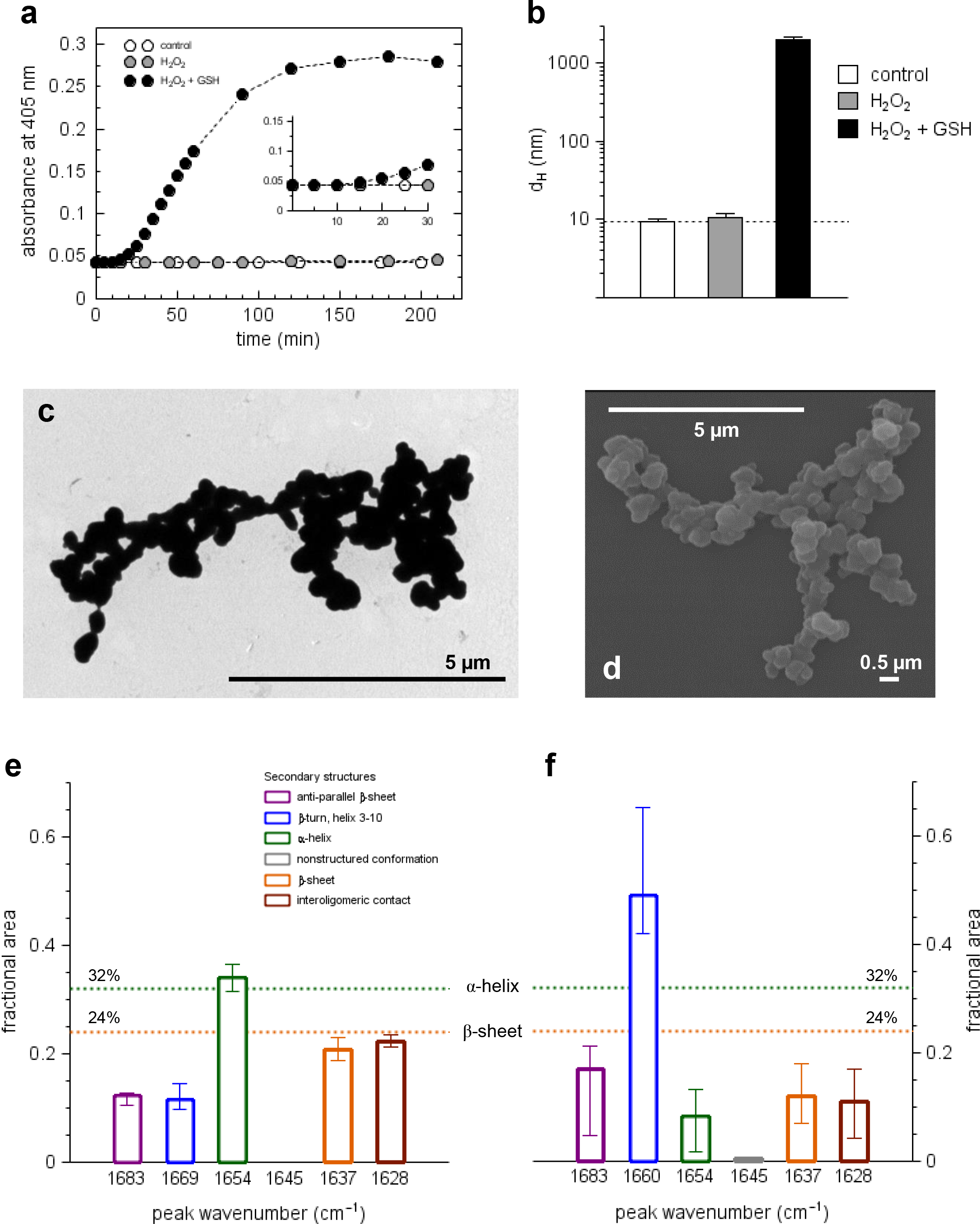
Oxidative treatments alter AtGAPC1 stability inducing globular aggregation. **(a)** Turbidity analyses monitored at Abs405 of AtGAPC1 (5 μM) incubated with H_2_O_2_ in the absence (grey closed circles) or presence of GSH (black closed circles). In the control experiment, change in turbidity was measured following incubation of AtGAPC1 in buffer alone (open circles). The turbidity of control and H_2_O_2_-treated AtGAPC1 showed no variation over 90 min while glutathionylated AtGAPC1 has a lag phase (15-20 min) followed by a rapid increase reaching a plateau after 2h incubation. Data correspond to the mean of three biological experiments. **(b)** Dynamic light scattering (DLS) measurement of AtGAPC1 incubated for 90 min in the presence of buffer alone (white bar), H_2_O_2_ (grey bar), or H_2_O_2_ supplemented with GSH (black bar). No appreciable variation of protein diameter (dH) was observed for control and H_2_O_2_-treated AtGAPC1 whilst glutathionylated AtGAPC1 formed aggregates with a diameter of ~2 μm. This value might be underestimated due to technical limitations of the DLS instrument linked to the polydispersity index of samples. Data represent mean ± s.d., *n* = 3 experiments with technical duplicates; \*\**P* < 0.01. Representative TEM **(c)** and SEM **(d)** images of aggregated AtGAPC1 obtained after 90 min incubation in the presence of H_2_O_2_ and GSH. (Scale bars: 5 μm and 0.5 μm). **(e)** Relative content in secondary structures of native AtGAPC1 determined by FTIR analysis. FTIR spectra were acquired after H_2_O to D_2_O substitution achieved through exhaustive concentrating / diluting steps. **(f)** Relative content in secondary structures of aggregated AtGAPC1 determined by FTIR analysis. The protein sample was incubated for 90 min in the presence of H_2_O_2_ and GSH and after treatment, the sample was centrifuged and the pellet resuspended in D_2_O. In panels **(e)** and **(f)**, the percentages of secondary structures derived from the crystallographic 3D-structure of native AtGAPC1 are also indicated as dashed lines.

The formation of products with increasing size was confirmed by dynamic light scattering (DLS) measurements, from which information about the size of the aggregates may be derived from autocorrelation functions. By fitting the autocorrelation function to a spherical model, a hydrodynamic diameter (d_H_) of 9.2 ± 0.5 nm was calculated for soluble AtGAPC1 (**Fig. 1b**). This value did not change over time and upon treatment with H_2_O_2_ alone (**Fig. 1b**), and was compatible with the crystal structure of AtGAPC1 tetramers^45^. On the other hand, aggregates formed after 90 min incubation with H_2_O_2_ and GSH were ~200-fold larger in terms of hydrodynamic diameter (**Fig. 1b**), thus roughly corresponding to the volume of 10 million AtGAPC1 tetramers.

Protein aggregates could be dissolved by SDS at 95 °C and migrated in reducing polyacrylamide gels as a single band corresponding to AtGAPC1 monomers (**Fig. S1a**). Under non-reducing conditions, the protein profile in SDS-PAGE was almost identical, except for a minor band likely corresponding to AtGAPC1 tetramers (**Fig. S1a**).

Inspection of AtGAPC1 aggregates by transmission and scanning electron microscopy (TEM and SEM, respectively) revealed irregular shapes resulting from the random binding of nearly globular particles of ~300-500 nm (**Fig. 1c** and **Fig. 1d**). No fibrils were observed. Fluorescence spectroscopy demonstrated that AtGAPC1 aggregates could interact with dyes commonly used to stain beta-enriched regions (Thioflavin-T, ThT) and hydrophobic patches (1-Anilino-8-Naphthalene Sulfonate, ANS) in protein aggregates (**Fig. S1b** and **Fig. S1c**, respectively). Native AtGAPC1, on the other hand, showed minimal interaction with either ThT or ANS dyes, suggesting that the conformation of AtGAPC1 in the aggregates was different from the native one.

A detailed analysis of secondary structure elements in both native and aggregated AtGAPC1 was performed by Fourier-transform infrared spectroscopy (FTIR). In order to test the method, the amide I band of native AtGAPC1 was decomposed into five Gaussian curves corresponding to different secondary structure motifs according to the literature^46, 47^ (**Fig. S2a**). The relative content of α-helices (1654 cm^−1^) and β-sheets (1637 cm^−1^) was in good agreement with crystallographic data (**Fig. 1e**). Additionally, the absence of a Gaussian component at 1645 cm^−1^ was fully consistent with the absence of disordered regions in native AtGAPC1 structure. The same measurements performed on AtGAPC1 aggregates showed an amide I band of different shape (**Fig. S2c**). The amide I difference spectrum between the two forms (**Fig. S2b**) exhibited an increase of absorption at wavenumbers higher than 1665 cm^−1^, paralleled by a decrease between 1665 and 1625 cm^−1^. The decomposition of the amide I band showed two major effects consisting in a 3-fold decrease of α-helices and a 4-fold increase in short structural motifs β-turns and 3_10_-helices; 1669 and 1660 cm^−1^ in native and aggregated protein, respectively), suggesting a conversion of the former into the latter ones (**Fig. 1f**). No clear changes were observed in other spectral components including unstructured regions (1645 cm^−1^) and β-sheets (1636 cm^−1^), the latter result suggesting that AtGAPC1 aggregates do not contain the cross-β spine typical of amyloid-like fibrils^48, 49^.

### 2. Aggregating AtGAPC1 is transiently glutathionylated

The treatment inducing AtGAPC1 aggregation (*i.e.* 125 μM H_2_O_2_ plus 625 μM GSH) also determined a rapid and complete inactivation of protein activity (**Fig. 2a**). Complete enzyme inactivation was attained in about 15 min and during this time, the inhibition of AtGAPC1 could be efficiently reverted by the thiol reducing agent dithiothreitol (DTT) (**Fig. 2b**). However, DTT became less and less effective over time and after 90 min incubation with H_2_O_2_ and GSH, only 25% of AtGAPC1 activity could be recovered by the reductive treatment (**Fig. 2b**).

**Figure 2.**
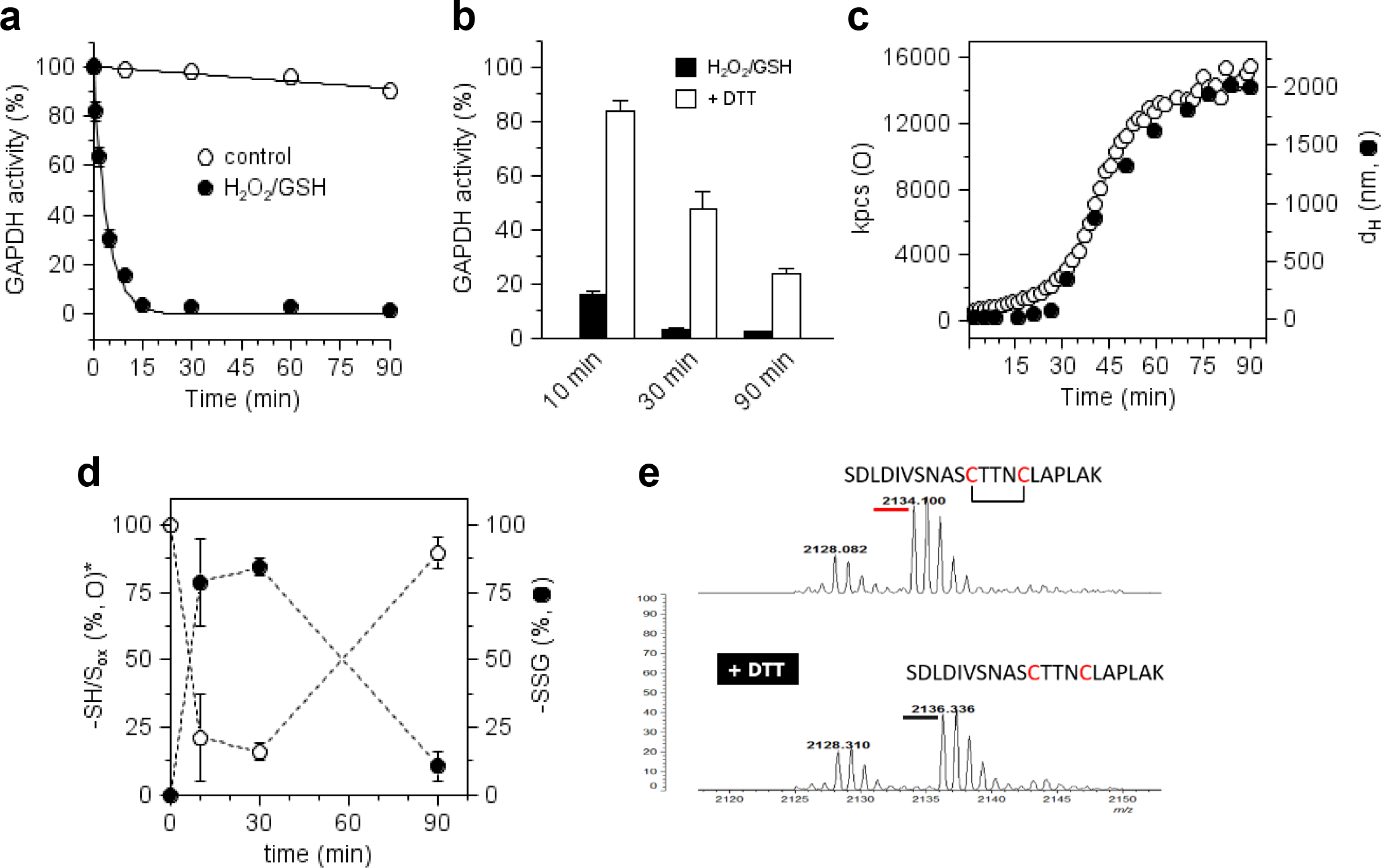
AtGAPC1 aggregation is specifically induced by transient S-glutathionylation. **(a)** Inactivation kinetics of AtGAPC1 in the presence of H_2_O_2_ and GSH. Treatment of AtGAPC1 with H_2_O_2_ and GSH causes a rapid inactivation with a complete loss of protein activity after 15 min. Data represent me an ± s.d., *n* = 3 experiments with technical duplicates. **(b)** Time-course DTT-dependent reactivity of inactivated AtGAPC1. At different time points, AtGAPC1 samples treated with H_2_O_2_ and GSH were further incuba ted with DTT to assess the inactivation reversibility, which was found to be strictly dependent on incubation times. Data represent mean ± s.d., *n* = 3 experiments with technical duplicates. **(c)** Time-course DLS analysis of AtGAPC1 tre ated with H_2_O_2_ and GSH. Protein diameters (dH, closed bla ck circles) and integrated count rates (kp cs, open circles) were monitored over time and plotted *versus* time (0-90 min). Data correspond to the mean of three biological experiments. **(d)** Time-dependent M ALDI-TOF signals of AtGAPC1 treated with H_2_O_2_ and GSH. Percentage of glutathionylated (*i.e.* ~300 Da shifted, closed circles) *versus* native (SH) or oxidized other than glutathionylated (S_ox_) (op en circles) AtGAPC1 forms were extrapolated from MALDI-TOF spectra and plotted *versus* times (0, 10, 30, and 90 min). Data represent mean ± s.d. (*n* = 3). **(e)** Peptide analysis of a ggreg ated AtGAPC1. Following 90 min treatment with H_2_O_2_ and GSH, the protein was centrifuged and the pellet was resuspended, subjected to tryptic digestion and analysed by mass spectrometry analysis prior and after treatment with DTT.

A detailed analysis by DLS, with light scattered intensity measured over 100 seconds intervals during the whole 90 min experiment, provided further hints on the aggregation process that paralleled the progressive, irreversible inactivation of AtGAPC1. The autocorrelation function, averaged over 500 seconds intervals, clearly demonstrated a slow increase of particle sizes during the first 25 min of the experiment (**Fig. 2c** and **Fig S3a**) followed by rapid growth in the micrometric range till the end of the experiment (**Fig. 2c**). The integrated count rate (kpcs), which depends on both the concentration and the size of the aggregates, showed a similar profile (**Fig. 2c**). Combination of the plots (**Fig. 2c**) thus suggests that AtGAPC1 is first induced to self-assemble into small nanoxparticles that only later start to melt into larger micrometric aggregates.

MALDI-TOF mass spectrometry (MS) analysis showed that AtGAPC1 was transiently glutathionylated during the aggregation process. A large portion of AtGAPC1 (~75%) became glutathionylated after only 10 min of treatment with H_2_O_2_ and GSH (1 GSH/ monomer, **Fig. 2d** and **Fig. S3b**) and reached ~85% after 30 min incubation (**Fig. 2d** and **Fig. S3b**). However, after 90 min incubation, glutathionylation decreased to roughly 10% (**Fig. 2d** and **Fig. S3b**) and dropped to zero if the MS analysis was carried out on the insoluble aggregates obtained by centrifugation (**Fig. S3c**).

In principle, the sulfur atom of GSH could have been attacked by either a water molecule or by a thiol. In the latter case, the nucleophilic thiol could either belong to a second GSH molecule or to Cys153, the only other cysteine of AtGAPC1. In order to solve the query, tryptic peptides were obtained from AtGAPC1 aggregates and analyzed by MALDI-TOF MS. The presence of an intramolecular disulfide bond between Cys149 and Cys153 (**Fig. 2e**) clearly demonstrated that the removal of GSH was the consequence of the nucleophilic attack performed by Cys153 of the same subunit on the proximal sulfur atom of glutathionylated Cys149.

Overall, the plots of **Figure 2**(panel **a-d**) hence describe a scenario in which AtGAPC1 is rapidly and reversibly inactivated by glutathionylation of Cys149 and slowly induced to self-assemble into small nanoparticles that later rapidly melt into larger micrometric aggregates. During this massive aggregation phase, inhibition of enzyme activity becomes permanent (not reverted by DTT) and glutathionylation of Cys149 is progressively substituted by a Cys149-Cys153 disulfide bond.

### 3. Structural snapshots along the pathway to AtGAPC1 aggregation

With the aim of describing the early steps of protein aggregation at the structural level, the crystal structure of AtGAPC1 was determined after treatments with H_2_O_2_ alone or H_2_O_2_ plus GSH.

Inhibition of AtGAPC1 activity by H_2_O_2_ alone is even faster than under glutathionylating conditions (H_2_O_2_ plus GSH) (**Fig. S4a** and **Fig. 2**), and is irreversible (**Fig. S4b**). This fast and irreversible kinetics indicates that the first oxidation product of catalytic Cys149 (sulfenic acid) is short-lived because it further reacts with H_2_O_2_ to generate more oxidized forms (sulfinic or sulfonic acids). Fully oxidized AtGAPC1 remains, however, fully soluble and tetrameric (**Fig. 1a**, **Fig. 1b**, and **Fig. S4c**).

After soaking AtGAPC1 crystals with H_2_O_2_, the effect of the oxidant could be directly evaluated from X-ray diffraction analysis and model structure calculations. The computed electron density map of H_2_O_2_-oxidized AtGapC1 showed a clear positive density in the *F*_o_−*F*_c_ map around the thiol group of catalytic Cys149. In this additional electron density, a sulfinic acid or sulfinate (−SO_2_H or −SO_2_^−^) was easily built (**Fig. S4d**), in agreement with recent quantum-mechanical analyses^45^. Interestingly, no additional positive electron densities were observed for other H_2_O_2_-sensitive residues, *i.e.* neither for the second cysteine (Cys153) nor for the seven methionines of each subunit (**Fig. S5**). This observation demonstrates the absolute selectivity of H_2_O_2_ for catalytic Cys149 among all other residues of AtGAPC1 in these conditions.

Crystals of AtGAPC1 were also soaked with a solution containing both H_2_O_2_ and GSH (1:10 ratio) with the aim of attempting a glutathionylation reaction *in crystallo*. Again, soaked crystals were subjected to X-ray diffraction analysis and the 3D structure was solved at 3.0 Å resolution (**Fig. 3a**). The calculated *F*_o_ − *F*_c_ electron density map revealed, in both chains O and R of the asymmetric unit, an elongated positive electron density stemming from the thiol group of Cys149, which could be interpreted as a mixed disulfide bond with a glutathione molecule (**Fig. 3b**). Additional, discontinuous, electron density regions were ascribed to the carboxylic groups of bound GSH (**Fig. 3b**). Beyond the mixed disulfide bond, GSH was stabilized by hydrogen bonds with the protein, the NAD^+^ cofactor and the sulfate ions bound to the active site (P_S_ and P_i_ sites)35 (**Fig. 3b**). Consistent with the results with H_2_O_2_-soaked crystals, no other residues than Cys149 were modified by the treatment with GSH and H_2_O_2_. Therefore, even in the crystal, AtGAPC1 may undergo the specific glutathionylation of Cys149 in the presence of H_2_O_2_ and GSH indicating that, under these conditions, sulfenic Cys149 (formed by the reaction of the thiol with H_2_O_2_) reacts faster with GSH than with H_2_O_2_, thereby escaping the irreversible modification.

**Figure 3.**
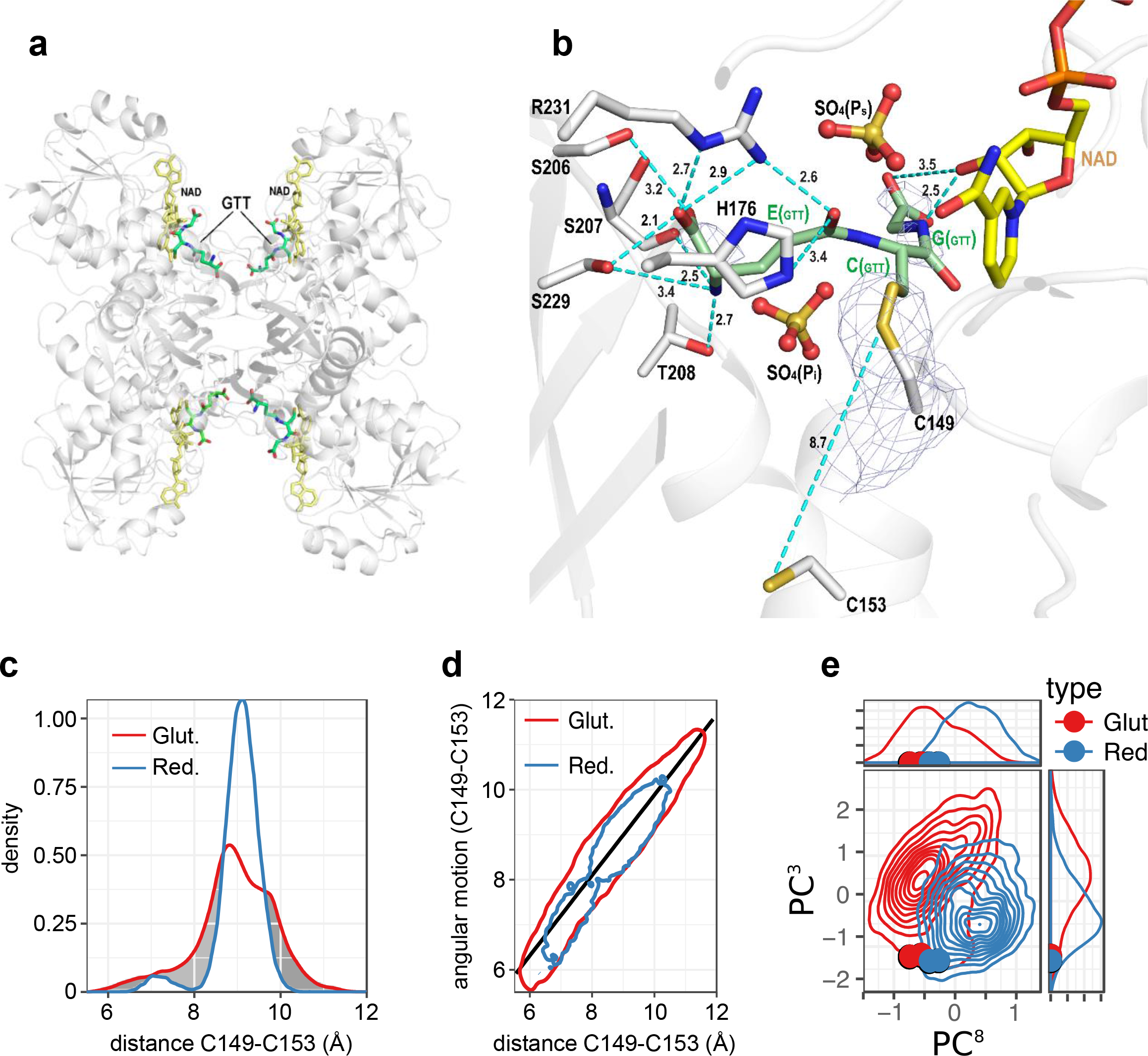
Structural features and MD simulation of S-glutathionylated AtGAPC1. **(a)** Cartoon representation of AtGAPC1 tetramer structure with four glutathione (GTTin the figure) molecules covalently bound to the catalytic cysteines and four c of actor molecules (NAD^+^) non-covalently interacting with the protein. Glutathione (GTT) and NAD^+^ molecules are shown as sticks. **(b)** Representation of the mixed disulfide bond between the catalytic cysteine (Cys149) and glutathione. The hydrogen bonds between glutathione and the protein (cut-off distance 3.5 Å) as well as the 2*F*_o_ − *F*_c_ electron density map (contoured at 1.5 σ) around the Cys149 and glutathione, are shown. The distance between the sulfur atoms of Cys149 and 153 is only slightly shorter in glutathionylated AtGAPC1 with respect to the unmodified protein (**Fig. S3d**). The interactions between GSH and sulfate ions (P_s_ and P_i_ sites) were omitted to mimic the solution environment **(c)** MD simulation-based analysis of distance distributions between the two sulfur atoms of Cys149 and Cys153 in reduced and glutathionylated forms are shown in blue and red colour, respectively. The distributions highlight the impact of glutathionylation on the cysteines 149-153 distance. **(d)** Correlation of a metric combining several key angles with the distance between the two sulfur atoms of cysteines 149 and 153, contour of the lowest density for the glutathionylated and reduced forms are indicated by a red and blue contour, respectively. **(e)** Principal component analysis (PCA) applied to the whole MD simulation dataset of GAPC1 in reduced and glutathionylated form. Projection of third (y-axis) and eighth (x-axis) PCA modes computed on protein secondary structure regions. An overall density contour plot for the whole simulation set consolidating the data on all four chains is displayed as blue and red plain lines for reduced and glutathionylated forms, respectively. Crystal structure principal component projections are indicated as coloured dots. The probability distribution of both PCA modes is shown above and to the right of the corresponding contour plot axes, respectively. Crystal structure projection on both modes are indicated as coloured dots.

Each GAPDH tetramer can bind a maximum of four GSH molecules lying at the entrance of the four active sites (**Fig. 3a**). However, GSH molecules did not occupy all the available sites in AtGAPC1 crystals and were characterized by high thermal parameters. Indeed, the GSH occupancy (q) was 82% for chain O and 67% for chain R, and thermal parameters (B) ranged between 60 and 80 Å^2^, indicating substantial mobility of the GSH molecules within AtGAPC1 active sites.

Multiple molecular dynamics (MD) simulations, starting from the crystal structure of glutathionylated AtGAPC1 confirmed the high mobility of bound GSH. Six main conformational clusters of the glutathione were observed (**Fig. S6**) with only one (cluster 2) corresponding closely to the starting crystal structure. Glutathionylation of Cys149 had no significant effect on the overall conformation of AtGAPC1 tetramers within the crystal (0.45 Å rmsd on C_α_ atoms between glutathionylated and native AtGAPC1) and the average distance between the S atoms of Cys149 and Cys153 remained prohibitive for the formation of a disulfide (8.7 Å; **Fig. 3b**). This result is consistent with the fact that the protein was crystallized before undergoing glutathionylation.

MD simulations indicated, however, that glutathionylation had an impact on the inter-sulfur distance between Cys149 and Cys153. The modification clearly extends the range of inter-sulfur distances, sampling both shorter and longer distances compared to the 8.8 Å of the native /reduced form (**Fig. 3c and Fig. S3d**). A strong correlation between the inter-cysteine distances and a metric combining key dihedral angles included between Cys149 and Cy153 was found (**Fig. 3d**), with a marked influence of the redox state. The overall conformational distribution of secondary structure elements analyzed by principal component analysis (PCA) showed that glutathionylation of Cys149 modified the protein conformational landscape and its plasticity by inducing a change in inter-domain arrangement (**Fig. 3e**). These changes observed on a microsecond timescale may represent the early stages of a local unfolding process that selectively prompted the protein to evolve towards an aggregation-prone conformation. Because of the subtle nature of the effects and the restricted timescale that can be sampled, quantitatively linking our observations to the formation of the Cys149-Cys153 disulfide and to the aggregation process on a much longer timescale remains, however, an important challenge.

### 4. The relevance of the Cys149-Cys153 disulfide bond for aggregation

The relevance of the Cys149-Cys153 disulfide bond for aggregation of AtGAPC1 was tested by chemical and genetic modifications. Preventing disulfide formation by mutating Cys149 into Ser had the effect of inhibiting both AtGAPC1 activity^45^ and its aggregation propensity in the presence of H_2_O_2_ and GSH (**Fig. 4a**). A similar effect was obtained when AtGAPC1 was pre-treated with the cysteine-alkylating agent iodoacetamide (IAM; **Fig. 4a**). Since IAM specifically alkylates Cys149^37^, this result confirms once again that aggregation is triggered by glutathionylation of Cys149, but provides no information on the role of the Cys149-Cys153 disulfide bond in the aggregation process. Mutation of Cys153 into Ser had, in contrast, no effect on enzyme activity and reversible inhibition by H_2_O_2_ and GSH (**Fig. S7a** and **Fig. S7b**). The C153S mutant was extensively glutathionylated after a 30 min incubation with H_2_O_2_ and GSH (**Fig. 4b** and **Fig. S7c**) and, in sharp contrast to wild type AtGAPC1, glutathionylation was stable over time, confirming the role of Cys153 in deglutathionylation (**Fig. 4b** and **Fig. S7c**). Interestingly, turbidity measurements showed that the aggregation kinetics of mutant C153S was 3.5-fold slower than that of wild type AtGAPC1 (**Fig. S7d**), in spite of the lag phase of similar duration (**Fig. 4c** and **Fig. 4d**). DLS analysis showed that both count rate increase (kcps; **Fig. 4e**) and particle size increase (d_H_; **Fig. 4f**) were significantly lower in the mutant compared to the wild type protein. Clearly, impairing the formation of the Cys149-Cys153 disulfide bond had a negative effect on the aggregation process. Conversely, the formation of the Cys149-Cys153 disulfide bond sped up the process and promoted the growth of larger aggregates.

**Figure 4.**
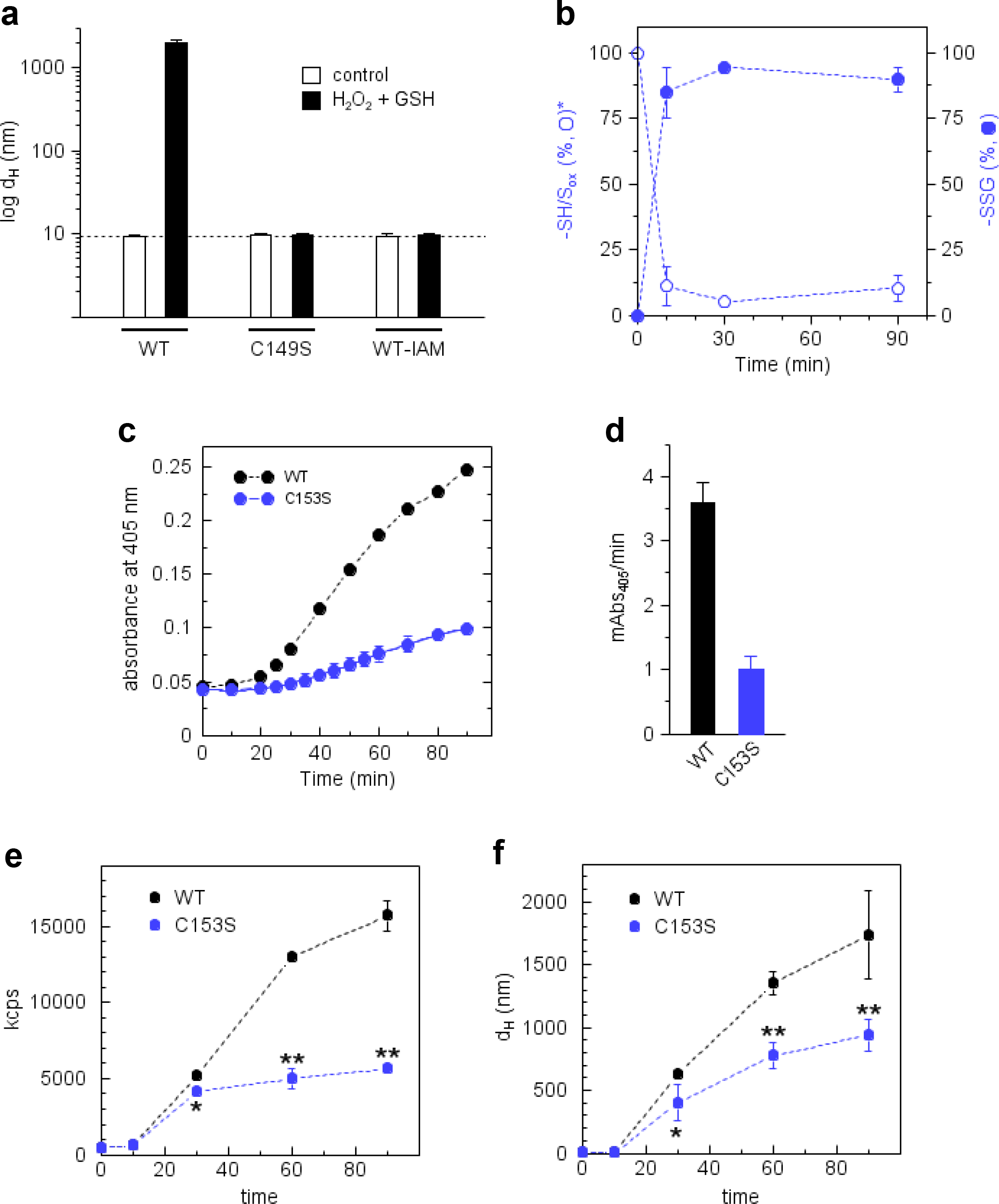
Effect of Cys153 and Cys149 on AtGAPC1 aggregation process. **(a)** DLS analysis of aggregation state of AtGAPC1 WT (WT), C149S mutant, and IAM-pretreated AtGAPC1 WT (WT-IAM) following 90 min incubation in the presence of buffer alone or H_2_O_2_ supplemented with GSH. While AtGAPC1 WT increased in size (~2 μM diameter), neither C149S mutant nor IAM-treated WT underwent an alteration of their tetrameric folding state. Data represent mean ± s.d., *n* = 3 experiments with technical duplicates. **(b)** Time-dependent MALDI-TOF signals of C153S treated with H_2_O_2_ and GSH. Percentage of glutathionylated (*i.e.* ~300 Da shifted, closed blue circles) *versus* native (not shifted, open blue circles) C153S forms were extrapolated from MALDI-TOF spectra and plotted *versus* times (0, 10, 30, and 90 min). Data represent mean ± s.d. (*n* = 3). **(c)** Turbidity analyses of AtGAPC1 WT and C153S mutant incubated with H_2_O_2_ and GSH. Compared to WT, the turbidity of C153S showed a similar lag phase (15-20 min) but a much slower increase. Data represent mean ± s.d., *n* = 3 experiments with technical duplicates. **(d)** Turbidity kinetics of AtGAPC1 WT and C153S (black and blue bar, respectively) expressed as milliA bs405 / min as derived by panel c. Data represent mean ± s.d., *n* = 3 experiments with technical duplicates. **(e-f)** Time-dependent DLS analysis of AtGAPC1 WT and C153S mutant treated with H_2_O_2_ and GSH. Integrated count rates (kcps) and protein diameter (dH) of AtGAPC1 WT (closed black circle) and C153S (closed blue circles) are shown in the panel **(e)** and **(f)**, respectively. Data represent mean ± s.d., *n* = 3 experiments with technical duplicates.

## DISCUSSION

In this work, we describe a mechanism by which an abundant tetrameric protein like GAPC1 from *Arabidopsis thaliana* is primed to form insoluble oligomeric aggregates (protein particulates) by the specific glutathionylation of its catalytic cysteine.

Due to its acidic and nucleophilic catalytic cysteine (Cys149), AtGAPC1 reacts quickly with H_2_O_2_. Cys149 thiolate (−S^−^) is first converted to a sulfenic acid intermediate (−SOH) and then to sulfinic acid (−SO_2_^−^)^45^. The modification of Cys149 by H_2_O_2_ is specific and irreversibly inhibits enzyme activity. The only other cysteine of AtGAPC1 (Cys153), which is found strictly conserved in GAPDH enzymes from almost all living organisms^41^, is less exposed to the solvent and remains reduced even after long incubations with H_2_O_2_. This residue, which is 8.8 Å apart from Cys149 in terms of inter-sulfur distance, does not react with sulfenic Cys149 either. *In vitro*, the two-step oxidation of Cys149 thiolate to the sulfinic acid proceeds undisturbed even at AtGAPC1:H_2_O_2_ equimolar concentrations^45^. *In vivo*, however, in the presence of millimolar concentrations of glutathione^50^, AtGAPC1 in Arabidopsis^51^, and orthologues in other photosynthetic organisms (*Synechocystis* sp. PCC6803^*52*^; *Chlamydomonas reinhardtii^53^*) are found glutathionylated under oxidative stress conditions. This observation indicates that sulfenic Cys149 may react faster with GSH than with H_2_O_2_. Efficient interference of GSH on AtGAPC1 irreversible oxidation apparently relies on the easy accommodation of glutathione within AtGAPC1 active sites where it can be stabilized by several interactions with the protein, as derived from our crystal structure of glutathionylated AtGAPC1 (**Fig. 3a** and **Fig. 3b**). From its preferential position, GSH may attack sulfenic Cys149 leaving no possibility for H_2_O_2_ to compete. As long as the activity of glutathionylated GAPC can be recovered by cytoplasmic glutaredoxins or thioredoxins^37^, the resulting glutathionylation/ deglutathionylation cycle perfectly fits into the metabolic remodelling of aerobic cells under oxidative stress conditions. Observations made on different types of aerobic cells show indeed that (i) H_2_O_2_ may inhibit glycolysis by inactivating GAPDH; (ii) the cell antioxidative response consumes NADPH for protective (recycling) functions including GAPDH deglutathionylation; and (iii) the pentose phosphate pathway and, only in plants, non-phosphorylating GAPDH, both activated by low NADPH levels, provide the NADPH required^35, 41, 44, 54^.

Although proposed as a salvage pathway for redox-sensitive GAPDH, here we show that glutathionylation destabilizes AtGAPC1 conformation and, in the long run, promotes the formation of insoluble aggregates.

Aggregation of AtGAPC1 develops as a three-phase process. Glutathione is quite mobile when it is covalently bound to Cys149, but it interacts with the protein and affects its dynamics, both globally and locally, as clearly shown by MD simulations (**Fig. 3d** and **Fig. 3e**). These changes are indicative of a conformational evolution whose effects need tens of minutes to show up. In our hands, AtGAPC1 could stay glutathionylated for about 10 min without changing significantly its overall native conformation. In this “pre-aggregation phase”, enzyme activity can be fully recovered upon removal of GSH by reduction (glutathionylation/deglutathionylation cycle)^38^.

True aggregation starts only later, during the “oligomeric phase”. In this phase, the DTT efficiency in recovering the AtGAPC1 activity starts to decline, indicating that oligomerization is associated with a permanently inactivated state of the protein. In principle, this effect may suggest that the protein is misfolded and no more active, or alternatively, that AtGAPC1 active sites are inaccessible to DTT because of initial aggregation. The “oligomeric phase” ends up with the completion of the lag phase of the whole aggregation process.

In the third phase, much faster than the previous one, particles abruptly start to grow to reach micrometric dimensions. Final aggregates are made by hundreds of bead-like particles of roughly 500 nm in d_H_ (≈ 10^5^ tetramers). Particle units are linked together to form irregular branched chains. During this “particulate phase”, AtGAPC1 loses its glutathionyl moiety in favour of a Cys149-Cys153 disulfide. This is a hallmark of an initial conformational shift. Otherwise, the formation of the Cys149-Cys153 disulfide would be prevented by the long inter-sulfur distance and unsuitable orientation of the side chains of the two cysteines that, in native AtGAPC1, are fixed into the rigid structure of a α helix (**Fig. S3d**). Once formed, the disulfide must be instrumental for aggregation, as suggested by the slower and limited aggregation of the C153S mutant. The conformational modification undergone by aggregated AtGAPC1 is documented by the (limited) change in secondary structures revealed by FTIR (**Fig. 1e, Fig. 1f**, and **Fig. S2**) and by the increased exposition of hydrophobic regions that bind ANS (**Fig. S1c**). The binding of ThT (**Fig. S1b**), documented by an increase in fluorescence that is orders of magnitude lower than that shown by amyloid fibrils^4^, is in agreement with the absence of a FTIR positive signal ascribable to inter-chain β-structures (1620 cm^−1^), thus excluding the possibility of a cross-β sheet architecture typical of amyloids. On the other hand, the lack of unstructured regions (1645 cm^−1^) and the maintenance of secondary structures overall, seem to exclude the possibility that aggregates are fully disordered and amorphous.

Shape and features of AtGAPC1 aggregates are reminiscent of animal GAPDH aggregates obtained after non-specific oxidation by the nitric oxide donor NOR3^21–23^, although more than 15 different residues of GAPDH, including cysteines, methionines, and tyrosines, were found modified by the NOR3 treatment^21^. Aggregates contained inter-molecular disulfides that bound together a minor portion of GAPDH subunits^21, 22^. These aggregates, grown *in vitro* following a nucleated process under strongly oxidizing conditions, are thought to recapitulate the GAPDH aggregates observed *in vivo* in different pathological conditions (*e.g.* Alzheimer, Parkinson, ALS)^34, 55, 56^ and under oxidative stress^23^. Plants’ GAPC is also extremely sensitive to oxidative stress and a common target of redox post-translational modifications^35^, but to our knowledge, the only report of cytoplasmic GAPC aggregation deals with Arabidopsis plants treated with the flagellin fragment flg22^32^. The flg22 is a pathogen-associated molecular pattern that elicits basal immunity and ROS production and causes GAPC-GFP to coalesce into cytoplasmic fluorescent puncta that are indicative of protein aggregation.

Globular particulates like those formed by glutathionylated AtGAPC1 are also formed by other aggregation-prone proteins under conditions that promote aggregation^4, 6, 57, 58^. Often protein particulates are made of proteins only partially unfolded. As a rule, they do not contain cross β sheets, but the involvement of β sheets and hydrophobic interactions in the aggregate formation are suggested by ThT and ANS binding.

In conclusion, here we show that a common thiol-based post-translational modification that occurs under oxidative stress conditions and consists in the addition of a glutathionyl moiety to a nucleophilic cysteine previously oxidized by H_2_O_2_, can trigger the formation of insoluble protein aggregates. This effect was studied in AtGAPC1, the major glycolytic GAPDH isoform of *Arabidopsis thaliana*, and was found to primarily depend on two cysteines (Cys149, Cys153) that are highly conserved in GAPDHs of any source. The collapse of AtGAPC1 into insoluble particles can be efficiently counteracted by GSH removal, but if AtGAPC1 remains glutathionylated for too long, aggregation proceeds undisturbed until all the protein collapses into micrometric globular particles. The whole process primarily depends on catalytic Cys149 that needs to be first oxidized and then glutathionylated. The prompt oxidation of Cys149 by H_2_O_2_ is assisted by Cys153 in a mechanism that favours metabolic remodelling of oxidatively stressed yeast cells^41^. Interestingly, fully conserved Cys153 is shown here to assist also the aggregation process. Whether AtGAPC1 aggregates are dead-end products of oxidation that cells need to dispose of or whether the native protein might be recovered by disassembling the particulates remains an open question.

## ONLINE METHODS

### Reagents

Unless otherwise indicated, chemicals were from Sigma-Aldrich. H_2_O_2_ was quantified spectrophotometrically using a molar extinction coefficient at 240 nm of 43.6 M^−1^ cm^−1^.

### Expression and purification of recombinant GAPDH from *Arabidopsis thaliana*

pET28a(+) expression vectors carrying the coding sequence for wild-type Arabidopsis GAPC1 and cysteine variants (C153S and C149S mutants) were transformed in *E. coli* BL21DE3 (Invitrogen). A starter culture (LB containing 50 μg/ml kanamycin) was grown overnight at 37 °C. The volume was expanded to 1 liter, and the culture was incubated until it reached an OD_600_ of ~0.5. Expression was induced by adding IPTG at the final concentration of 0.1 mM. After 16-18 h of incubation at 25 °C, bacteria were pelleted by centrifugation (5,000 *g*, 15 min, 4 °C), resuspended in buffer A (30 mM Tris-HCl, pH 7.9, 100 mM NaCl and 5 mM imidazole) and lysed by sonication as described previously (Bedhomme et al., 2012 Biochem J). Lysates were centrifuged (30,000 *g*, 30 min, 4 °C) and supernatants passed through a 0.22 μm filter. Filtered supernatants were passed over a Ni^2+^ His-Trap chelating resin pre-equilibrated with buffer A. His-tagged proteins were then eluted according to the manufacturer’s instructions. Eluted fractions were analyzed by SDS-PAGE to assess purity grade. Selected fractions were pooled and desalted against buffer B (50 mM potassium phosphate, pH 7.5) using PD-10 columns (GE Healthcare). Desalted proteins were stored in buffer B at −20 °C. The concentration of purified proteins was determined spectrophotometrically using a molar extinction coefficient at 280 nm of 40,910 M^−1^ cm^−1^.

### GAPDH activity assay

The activity of GAPDH was monitored as described previously (Zaffagnini et al., 2007 FEBS J). Briefly, assays contained 20-50 nM GAPDH (WT and cysteine variants), 5 mM MgCl_2_, 3 mM 3-phosphoglycerate, 5 units ml^−1^ of baker’s yeast PGK, 2 mM ATP, 0.2 mM NADH and 1 mM EDTA in 50 mM Tris-HCl (pH 7.5). Initial rates were determined at 25 °C by measuring the decrease of absorbance at 340 nm (NADH oxidation) during the first 1-2 min using a Cary60 UV /Vis spectrophotometer (Agilent Technologies).

### Oxidation treatments of recombinant AtGAPC1

Recombinant proteins were incubated with 10 mM dithiothreitol (DTT) for 30 min and then desalted using NAP-5 columns (GE Healthcare) pre-equilibrated with 50 mM Tris-HCl, 1 mM EDTA, pH 7.5 (buffer C). For oxidation experiments, pre-reduced samples (5 μM) were treated with 0.125 mM H_2_O_2_ alone or supplemented with 0.625 mM GSH. All treatments were carried out at 25 °C in buffer C supplemented with 0.14 mM NAD^+^. At the indicated times, an aliquot was withdrawn to assay residual GAPDH activity. Activity data expressed as a percentage of maximal activity were plotted versus time. Interpolation curves were generated by non-linear regression using CoStat (CoHort Software). For recovery assays, 20 mM DTT were added at different time points to AtGAPC1 treated with H_2_O_2_ alone or supplemented with GSH. After 20 min incubation, aliquots were withdrawn for the assay of GAPDH activity.

### Aggregation kinetics measured by turbidity

Kinetics of AtGAPC1 aggregation were assessed by measuring the increase of turbidity at 405 nm. AtGAPC1 samples (WT and Cys variants) were incubated at 25 °C with or without 0.125 mM H_2_O_2_ alone or supplemented with 0.625 mM GSH in a low-protein-binding 96-well plate. All treatments were performed in buffer C supplemented with 0.14 mM NAD^+^. Samples were monitored over time, and turbidity at 405 nm was measured using a plate reader (Victor3 Multilabeling Counter; Perkin Elmer).

### Dynamic light scattering

Size distribution of the species present in samples were obtained from measurements of AtGAPC1 (WT and Cys variants) incubated with or without H_2_O_2_ alone or supplemented with GSH in buffer C plus 0.14 mM NAD^+^ at 25 °C. The data were obtained using Zetasizer Nano (Malvern) cuvettes and thirty spectra were acquired for each DLS analysis, averaged and used to determine the hydrodynamic diameter and polydispersity using the average autocorrelation function.

### ANS fluorescence assay

AtGAPC1 samples were incubated with or without H_2_O_2_ supplemented with GSH as described above. After 90 min incubation, aliquots were withdrawn and treated with 5 μM 1-anilinonaphthalene-8-sulfonic acid (ANS). Emission fluorescence spectra were acquired using a Cary Eclypse spectrofluorometer (Varian) in a 400-600 nm range at a 50 nm min^−1^ scan rate with an excitation wavelength of 380 nm and 5 nm slit width.

### Thioflavin-T fluorescence assay

AtGAPC1 samples were incubated with or without H_2_O_2_ supplemented with GSH as described above. After 90 min incubation, aliquots were withdrawn and treated with 25 μM thioflavin-T (Th-T). Emission fluorescence spectra were recorded after 2 min equilibration using a Cary Eclypse spectrofluorometer (Varian) in a 450-650 nm range at a 50 nm min^−1^ scan rate with an excitation wavelength of 445 nm and 5 nm slit width.

### Crystallization and data collection

Crystals of NAD^+^-AtGAPC1 were grown as previously reported in (Zaffagnini et al. 2016 ARS). Native crystals were soaked in the reservoir solution composed by 3.0 M (NH_4_)_2_SO_4_ and 0.1 M Hepes-NaOH (pH 7.5) plus 0.1 mM H_2_O_2_ to obtain oxidized AtGAPC1 crystals or plus 0.1 mM H_2_O_2_ and 1 mM GSH to obtain glutathionylated AtGAPC1 crystals. The H_2_O_2_/GSH molar ratio used in soaking experiments was doubled with respect to biochemical assays as the protein molecules were constrained in a crystalline lattice. Preliminary tests showed that the AtGAPC1 crystals were stable in the soaking solutions. The soaking was performed for 1 month then the crystals were fished and briefly soaked in a cryo solution containing 3.2 M (NH_4_)_2_SO_4_ and 20% v/v glycerol.

Diffraction data were collected using the synchrotron radiation of Elettra (Trieste, Italy) XRD1beam line, at 100 K with a wavelength of 1.26 Å, an oscillation angle of 1° and a sample-to-detector (Pilatus 2M) distance of 250 mm for oxidized AtGAPC1 and 260 mm for glutathionylated AtGAPC1 crystals. The images were indexed with XDS (Kabsch 2010, Acta Cryst. D.) and scaled with SCALA (Evans 2006, Acta Cryst. D) from the CCP4 package. The unit cell parameters and the data collection statistics are reported in Table S1.

### Structure Solution and Refinement

Since the unit cell parameters were not affected by the soaking, the AtGAPC1 structure (PDB code 4Z0H, Zaffagnini et al., 2016 ARS) was directly used in the refinement against the experimental data of the oxidized and glutathionylated AtGAPC1 crystals. After an energy minimization and B factor refinement with CNS1.3 (Brünger et al., 1998 Acta Cryst. D) selecting 5% of reflections for R_free_ calculation, the starting R and R_free_ were 0.289 and 0.361 for oxidized AtGACP1, and 0.237 and 0.287 for glutathionylated AtGAPC1. The calculated Fo-Fc electron density map clearly showed for both cases large positive electron densities in correspondence and near the thiol group of residue Cys149. In the case of oxidized AtGAPC1 a sulphinic group was built, while for glutathionylated AtGAPC1 a glutathione molecule forming a mixed disulphide bond with Cys149, was modeled into the densities. The manual rebuilding was performed with Coot (Emsley and Cowtan 2004 Acta Cryst. D). In the final stages of refinement performed with REFMAC5.5 (Murshudov et al., 2011 Acta Cryst. D) water molecules were automatically added, and after a visual inspection they were conserved in the model only if contoured at 1.0 σ on the (2*F*_o_ - *F*_c_) map and if they fell into an appropriate hydrogen bonding environment. Stereo-chemical quality of the models was checked with PROCHECK (Laskowski et al., 1993) and protein superimpositions were performed by LSQKAB (Vagin and Teplyakov 2010 Acta Cryst. D) from CCP4 package. Refinement statistics are reported in Table S1.

### Fourier transform infrared (FTIR) analysis

FTIR absorption measurements were performed at room temperature with a Jasco Fourier transform 6100 spectrometer, equipped with a DLATGS detector. The spectra were acquired with 2 cm^−1^ resolution in the whole mid-IR range (7000-1000 cm^−1^), using a standard high-intensity ceramic source and a Ge/KBr beam splitter. Ten μl of AtGAPC1 solution sample at 10 mg ml^−1^ in D_2_O was deposited between two CaF_2_ windows, separated by a 50 μm teflon spacer, and mounted into a Jasco MagCELL sample holder. All spectra were obtained by averaging 10^3^ interferograms.

H_2_O to D_2_O substitution was performed in the samples to avoid the overlapping of the spectral contribution due to the H_2_O bending mode, at around 1640 cm−1 (Marechal, 2011 J. Mol. Struct.), with the amide I band of the AtGAPC1, centered at approximately 1650 cm^−1^. Starting from 1 ml containing native AtGAPC1 (0.2 mg ml^−1^) in buffer C supplemented with 0.14 mM NAD+, D_2_O substitution was achieved by progressive dilution of H_2_O in D_2_O through several concentration-dilution cycles, by using Centricon Amicon filters with a cut-off of 10 kDa. The procedure was applied up to a final, nominal dilution corresponding to 1 H_2_O:7×10^4^ D_2_O. The efficacy of replacement was tested by comparing the FT-IR spectrum of the flow-through solution with that of pure D_2_O; the comparison clearly showed that in the spectral interval of the amide I (1750-1570 cm^−1^) the two spectra essentially coincided (not shown). The D_2_O-substituted AtGAPC1 solution was finally concentrated up to 10 mg ml^−1^. Spectral analysis of the amide I bands was performed after subtracting the spectrum of the buffer solution (*i.e.* the final flow-through of the D_2_O-substitution procedure), and by approximating the residual background with a cubic spline function with the OriginPro 9.1 software (OriginLab Corporation, Northampton MA, USA). Second and fourth derivative spectra in the amide I’ region were calculated using the *i-signal* program (version 2.72) included in the SPECTRUM suite (SPECTRUM: A Signal Processing Toolkit, Version 1.1, 1990) written in MATLAB language. A Savitsky-Golay algorithm was employed to smooth the signals and calculate the derivatives. The smooth width was chosen by evaluating step-by-step the impact of increasing the smoothing on the calculation of the derivative spectrum, with the aim of optimizing the signal to noise ratio without loosening spectral information. The decomposition of the amide I into Gaussian components was performed by using a locally developed least-squares minimization routine (Malferrari et al., 2015 J Phys Chem B), based on a modified grid search algorithm (Bevington, 1969). Confidence intervals for the best-fit parameters were evaluated numerically, as detailed in (Francia et al., 2009 J Phys Chem B). The peak wavenumbers of the Gaussian components were fixed to values inferred from the second and fourth derivative analysis, while the areas and the widths were treated as free parameters. Since both the second and the fourth derivative spectra can in principle exhibit artefactual peaks (*i.e.* not reflecting genuine components of the original spectrum) (Butler, 1970), we considered as reliable peaks only those which were present at the same wavenumber (within our spectral resolution, 2 cm^−1^) both in the second and in the fourth derivative spectra. This criterion resulted for the control in peaks fixed at 1683, 1669, 1654, 1637 and 1628 cm^−1^.

The aggregated AtGAPC1 was prepared by incubating the protein sample with H_2_O_2_ and GSH as described earlier. After 90 min incubation, the sample solution was centrifuged at 10,000 *g* for 15 min and the resulting pellet was resuspended with an adequate D_2_O volume to attain 10 mg ml^−1^ protein concentration. FT-IR analysis was carried out as described before and six Gaussian components with peak wavenumber fixed at 1682, 1660, 1652, 1644, 1636 and 1624 cm^−1^ were used.

### Electron microscopy

AtGAPC1 samples were incubated with H_2_O_2_ supplemented with GSH as described above, then deposited onto carbon-coated copper mesh grids, and negatively stained with 2% (w/v) uranyl acetate. The excess stain was wicked away, and the sample grids were allowed to air dry. The samples were viewed with an FEI Tecnai 12 BioTwin 85 kV transmission electron microscope (TEM), and digital images were taken with an Advanced Microscopy Techniques camera. The scanning electron microscope (SEM) observations were conducted using a Hitachi S-4000. The samples were deposited on a mica layer and gold coated (2 nm) prior the observations.

### MALDI-TOF Mass spectrometry

AtGAPC1 samples (WT and C153S mutant; 5 μM) were incubated at 25 °C with 0.125 mM H_2_O_2_ supplemented with 0.625 mM GSH in buffer C containing 0.14 mM NAD+. At different time points, aliquots were withdrawn for MALDI-TOF mass spectrometry analysis using a Performance Axima MALDI-TOF mass spectrometer (Shimadzu-Kratos, Manchester, United Kingdom) equipped with a 337 nm nitrogen laser. For mass determination of AtGAPC1 samples, spectra were acquired as described previously (Berger et al., 2016, Plant Physiol) with a pulse-extraction fixed at 60,000. For each time point, 4 μl of the reaction mixture were mixed with 2 μl of 1% trifluoroacetic acid (TFA) and then kept on ice to quench the reaction. To assess the effect of a reducing treatment, 4 μl of the reaction mixture was incubated for 15 min at 25 °C with 2 mM DTT in 50 mM ammonium bicarbonate before being processed as described above. Immediately after acidification, 1 μl of sample and 3 μl of a saturated solution of sinapinic acid in 30% acetonitrile containing 0.3% TFA were mixed together and 3 μl were deposited onto the sample plates for MS analysis. Spectra were acquired and processed as followed using the Launchpad software (Shimadzu-Kratos, version 2.8). Briefly, no baseline subtraction was performed but raw data were smoothed using an algorithm averaging MS data within a smoothing filter width of 40 time channels. Protein peaks were delimited manually and the software for both mass determination and peak area quantification used these limits. Peak areas were expressed as a percentage of the sum of all areas corresponding to AtGAPC1 proteoforms.

For peptide analysis, the aggregated AtGAPC1 was prepared by incubating the protein sample with H_2_O_2_ and GSH as described above in a 500 μl reaction mixture. After 90 min incubation, the sample was centrifuged for 5 min (10,000 *g*, 20 °C) and the resulting pellet was solubilized in 11 μl of 50 mM ammonium bicarbonate containing 5 mM iodoacetamide and 0.01% of MS-compatible ProteaseMax detergent (Promega). After resolubilization, 5 μl were mixed with 1 μl of a mixture of Trypsin/Lysine C endoproteases (Promega) prepared at 0.1 μg μl^−1^ in 1 mM HCl and leaved at 30 °C for 20 min. After incubation, 5 μl of 50 mM ammonium bicarbonate were added and digestion was completed at 30 °C. To assess the effect of reducing agents, one-half of the digestion mixture was subjected to DTT reduction. Tryptic digests were then acidified as described above and prepared for MALDI-TOF mass spectrometry as described in (Morisse et al., 2014 JBC). Peptide mass fingerprints were acquired in the positive reflectron ion mode after external calibration and processed with the same software as described above. Raw MS spectra were smoothed with a Gaussian algorithm using a smoothing filter width of 2 time channels. Baselines of MS spectra were also corrected using a baseline filter of 6 time channels. Monoisotopic peak recognition of peptides present in peptide mass fingerprints was done using the Poisson peptide algorithm and monoisotopic peaks were considered if they had at least 2 additional isotopes and showed with them a maximum intensity variation of 80%.

### Molecular dynamics

We set up simulation systems for AtGAPC1 in complex with NAD^+^ in both reduced and glutathionylated form using 3 different treatments of the NAD^+^ cofactor leading to 6 independent simulations. The first simulation set was run without any constraints. At the end of the production, for both reduced and glutathionylated forms, 3 out of 4 NAD^+^ left their binding sites. Although such partial occupation is not unexpected, we wanted to investigate here also the scenario of full occupation of all sites. For this purpose, we introduced distance restraints between NAD^+^ and protein C_α_ atoms in the two other simulation sets. In simulation set 2, distance restraints were applied between the two NAD^+^ ribose O4’ atoms and the closest Gly11 and Ile13 C_α_ atoms with a force constant of 500 kJ mol^−1^ nm^−2^. In simulation set 3 the same two distance restraints were applied with a higher force constant of 2000 kJ mol^−1^ nm^−2^ and two additional restraints involving the NAD^+^ adenine N3^’^ and N315 C_α_ atoms, and the NAD^+^ nicotinamide C4’ and D34 C_α_ atoms.

In all simulations, summarized in Table S2, Protein Data Bank ID 4Z0H was used as starting structure for the NAD^+^-GAPC1 complex in reduced form, whereas the crystal structure presented in this work was used for the glutathionylated form. Default protonation at pH 7.0 was used for all residues with the exception of the catalytic residues; His176 was doubly protonated and Cys149 was deprotonated for the reduced form or linked to the glutathione molecule in the glutathionylated form. Both forms were solvated in a dodecahedron box using periodic boundary conditions and a ~1.5 nm water shell. In four of the six simulation systems (numbered from 1 to 4 in Table S2) only a minimal number of counter ions was inserted to neutralize the charge of the protein (12 sodium ions). In the two remaining systems, ~150 mM of salt (Na^+^ Cl^−^) were included.

Molecular dynamics simulations were carried out using the CHARMM 36 force field for protein and the TIP3P model^4^ for water. The Gromacs 4.5 software was used to run the simulations using the virtual interaction sites approach, allowing a 5fs integration time step. Neighbor searching was performed every 5 steps. All bonds were constrained using the LINCS algorithm. The PME algorithm was used for electrostatic interactions with a cut-off of 1 nm. A single cut-off of 1 nm was used for Van der Waals interactions. Two independent baths for protein and solvent were coupled to a temperature of 310 K using the Bussy velocity rescaling thermostat with a time constant of τ = 0.1 ps. Pressure coupling was scaled isotropically using the Berendsen weak barostat to a reference pressure of 1 bar, τ = 5.0 ps and compressibility of 4.5 10^−5^ bar^−1^.

All systems were minimized for 10,000 steps with a steepest descent algorithm, equilibrated for 10 ns, using position restraints of 1000 kJ mol^−1^ nm^−2^ on heavy atoms with the crystal structure as a reference, followed by an equilibration of 20 ns using position restraints of 1000 kJ mol^−1^ nm^−2^ on C_α_ atoms, and finally 20 ns using position restraints of 10 kJ mol^−1^ nm^−2^ on C_α_ atoms. Production runs were computed for 1 μs without any position restraints.

### Accession Number

The atomic coordinates and structure factors of the oxidized AtGAPC1 (AtGAPC1-SO2H) and glutathionylated AtGAPC1 (AtGAPC1-SSG) have been deposited in the Protein Data Bank under accession codes PDB XXXX and PDB XXX, respectively.

---

Legends to Supplementary Figures

**Table S1:**
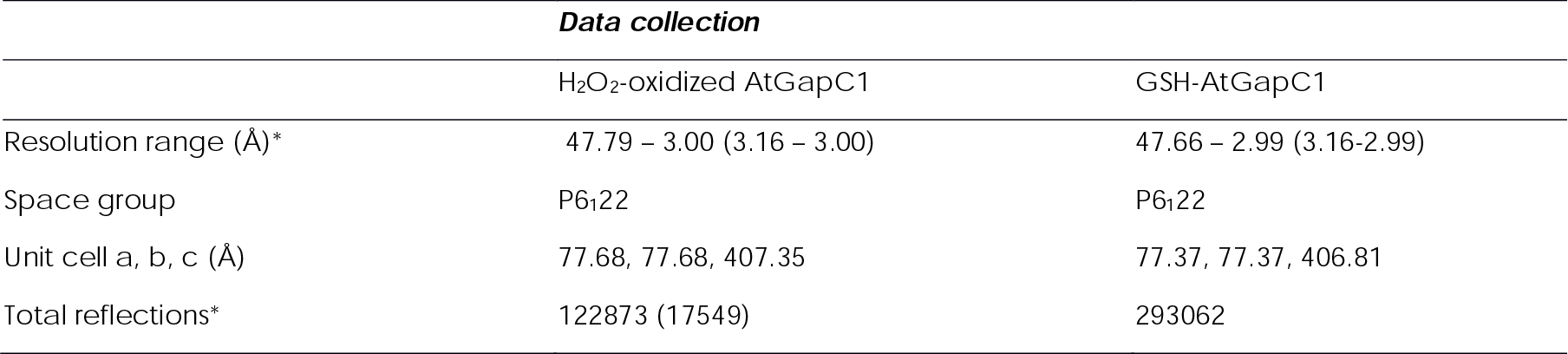
Data collection and refinement statistics

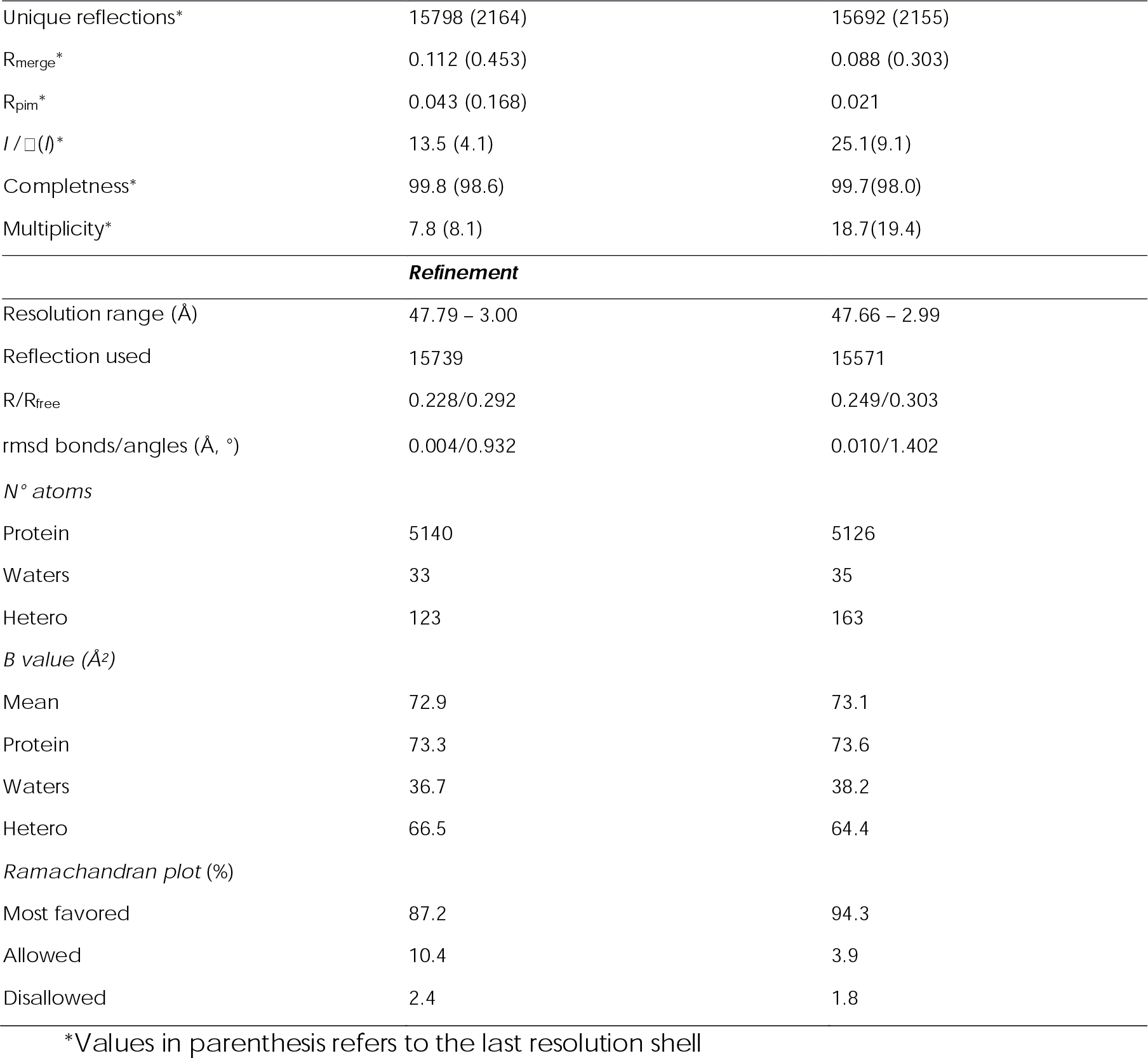

**Table S2:** Simulation systems overview

| Simulation Set | # | Acronym | Functional form | [Ion] (mM) | Constraint force on NAD<sup>+</sup> (kJ.mol<sup>-1</sup>.nm<sup>-2</sup>) | Composition (water/Na<sup>+</sup>/Cl<sup>-</sup>) | Atom number |
| --- | --- | --- | --- | --- | --- | --- | --- |
| 1 | 1 | Red | *reduced* | ~ 0 | 0 | 27033 / 12 / 0 | 104075 |
|  | 2 | Glut | *glutathionylated* | ~ 0 | 0 | 26741 / 12 / 0 | 103051 |
| 2 | 3 | Red_low_constr | *reduced* | ~ 0 | 500 | 27033 / 12 / 0 | 104075 |
|  | 4 | Glut_low_constr | *glutathionylated* | ~ 0 | 500 | 26741 / 12 / 0 | 103051 |
| 3 | 5 | Red_high_constr | *reduced* | ~ 150 | 2000 | 26889 / 84 / 72 | 103787 |
|  | 6 | Glut_high_constr | *glutathionylated* | ~ 150 | 2000 | 26597 / 84 / 72 | 102673 |

**Figure S1. (a)** AtGAPC1 samples (5◻DμM) incubated for 90 min in the presence of buffer (control) or 0.125 mM H_2_O_2_ supplemented with 0.625 mM GSH were loaded on a 4-20% SDS-PAGE under reducing (+) and non-reducing conditions (−). **(b)** Fluorescence emission spectrum of thioflavin T (ThT) with untreated (black line) or treated AtGAPC1 (red line). The treatment consisted of 90 min incubation of AtGAPC1 in the presence of buffer or H_2_O_2_ supplemented with GSH. Binding of amyloid-specific ThT to the aggregated AtGAPC1 resulted in a 2-fold increase of fluorescence emission at 482 nm. **(c)** Fluorescence emission spectrum of 1-anilinonaphthalene-8-sulfonic acid (ANS) with untreated (black line) or treated AtGAPC1 (red line). The treatment consisted of 90 min incubation of AtGAPC1 in the presence of buffer or H_2_O_2_ supplemented with GSH. Binding of ANS to the aggregated AtGAPC1 resulted in a red-shift of the peak and a concomitant ~10-fold increase of maximal fluorescence emission.

**Figure S2. FTIR analysis of native and aggregated AtGAPC1** FTIR spectra of the amide I region of native **(a)** and aggregated **(c)** AtGAPC1. The individual Gaussian components, corresponding to the different secondary structure elements, are plotted using the same colour code of Figure 1 (panel e and f), and the resulting fit is shown as a red curve (see “Online Methods” for details of the analysis). **(b)** Difference FTIR spectra between native and aggregated AtGAPC1.

**Figure S3. Effect of S-glutathionylation on AtGAPC1 (a)** Time-course DLS analysis of AtGAPC1 treated with H_2_O_2_ and GSH. The protein diameter (closed circles) was monitored as described in Figure 2c and plotted versus time (0-30 min). Data represent mean ± s.d., *n* = 3 experiments with technical duplicates. **(b)** MALDI-TOF spectra of AtGAPC1 treated with H_2_O_2_ and GSH. At indicated time points, AtGAPC1 samples were withdrawn, mixed in a ratio 2:1 (v/v) with sinapinic acid and analysed by MALDI-TOF MS to assess the redox state of the protein: glutathionylated (–SSG, ~305 Da shifted) *versus* native/oxidized (–S_ox_, not shifted). The peak highlighted by an asterisk corresponds to the protein-matrix adduct. The spectra are representative of three biological replicates. **(c)** Distance between the thiol groups of the catalytic Cys149 and the Cys153 in AtGAPC1 structure (PDB ID 4Z0H; Zaffagnini et al., 2016 ARS). The α-helix containing the two cysteine residues displayed as sticks is highlighted in yellow.

**Figure S4. Treatment with H_2_O_2_ does alter AtGAPC1 without affecting protein folding (a)** Inactivation kinetics of AtGAPC1 in the presence of H_2_O_2_. Treatment of AtGAPC1 with H_2_O_2_ causes a rapid inactivation with a complete loss of protein activity after 10 min. Data represent mean ± s.d., *n* = 3 experiments with technical duplicates. **(b)** Time-course DTT-dependent reactivity of inactivated AtGAPC1. At different time points, AtGAPC1 samples treated with H_2_O_2_ were further incubated with DTT to assess the inactivation reversibility, which was found null at each incubation times. Data represent mean ± s.d., *n* = 3 experiments with technical duplicates. **(c)** Size exclusion chromatography of native and H_2_O_2_-treated AtGAPC1. The protein maintained its tetrameric structure under both control (dotted line) and oxidizing (continuous line) conditions. Molecular masses of standard proteins are indicated as closed circles. **(d)** Representation of the crystal structure of oxidized AtGAPC1. The dimer (chains are named O and Q) composing the asymmetric unit is shown as cartoon and surface while the symmetry-related subunits completing the tetramer, are shown as a surface. The oxidized catalytic cysteines (sulphinic group) and the cofactor (NAD) are shown as sticks. A magnification of the catalytic cysteines with their environment and the 2*F*_o_ − *F*_c_ electron density map(contoured at 1.5 σ), is also reported.

**Figure S5. Position and accessibility of methionine residues and Cys153** The position of the seven methionine residues found in the sequence of AtGAPC1, and Cys153 is shown. The residues are represented as sticks. The values of calculated accessible surface area (ASA) are: Met40(O/Q) = 1.8/2.0 Å^2^; Met43(O/Q) = 10.8/10.6 Å^2^; Met127(O/Q) = 13.0/13.2 Å^2^; Met172(O/Q) = 0.1/0.2 Å^2^; Met190(O/Q) = 83.1/ 83.0 Å^2^; Met228(O/Q) = 5.6/5.9 Å^2^; Met328(O/Q) = 10.2/10.3 Å^2^; Cys153(O/Q) = 0.0/0.0 Å^2^.

**Figure S6. Molecular dynamics simulations on glutathionylated AtGAPC1** The main conformational clusters of the glutathione obtained from multiple molecular dynamics (MD) simulations (1 μs) starting from the crystal structure of glutathionylated AtGAPC1, are shown. The bottom graphs assign each simulation snapshot to one of the six conformations using a color code. Clusters confirmed the high mobility of bound GSH. Cluster 2 is the closest to the starting crystal structure. The glycine residue (G_GTT_) of glutathione is more mobile than the glutamate residue (E_GTT_) that forms several interactions with protein residues in most clusters (cut-off distance 3.5 Å). The structural clustering of glutathione conformations was achieved after computing the RMSD matrix between all pairs of glutathione coordinates, using the gromos algorithm (Angew. Chem. Int. Ed. 1999, 38, pp 236-240) with a cutoff of 2.5 Å, for each of the four chains of the three simulations of AtGAPC1 in the glutathionylated form.

**Figure S7. Effect of S-glutathionylation on the C153S mutant** (a) Inactivation kinetics of the C153S mutant in the presence of H_2_O_2_ and GSH. Treatment of C153S with H_2_O_2_ and GSH causes a rapid inactivation with a complete loss of protein activity after 15 min incubation. Inset, inactivation kinetics of AtGAPC1 WT and C153S revealed an identical response of both proteins to the inactivation treatment. Data represent mean ± s.d., *n* = 3 experiments with technical duplicates. **(b)** Time-course DTT-dependent reactivity of inactivated C153S. At different time points, C153S samples treated with H_2_O_2_ and GSH were further incubated with DTT to assess the inactivation reversibility, which was found to be strictly dependent on incubation times. Data represent mean ± s.d., *n* = 3 experiments with technical duplicates. **(c)** MALDI-TOF spectra of C153S mutant treated with H_2_O_2_ and GSH. At indicated time points, C153S samples were withdrawn, mixed in a ratio 2:1 (v/v) with sinapinic acid and analysed by MALDI-TOF to assess the redox state of the protein: glutathionylated (–SSG, ~305 Da shifted) *versus* native/oxidized (–S_ox_, not shifted). The peak highlighted by an asterisk corresponds to the protein-matrix adduct. The spectra are representative of three biological replicates.

