## Supplementary figures and images for "Glutathionylation primes soluble GAPDH for late collapse into insoluble aggregates"

### Supplemental Figures

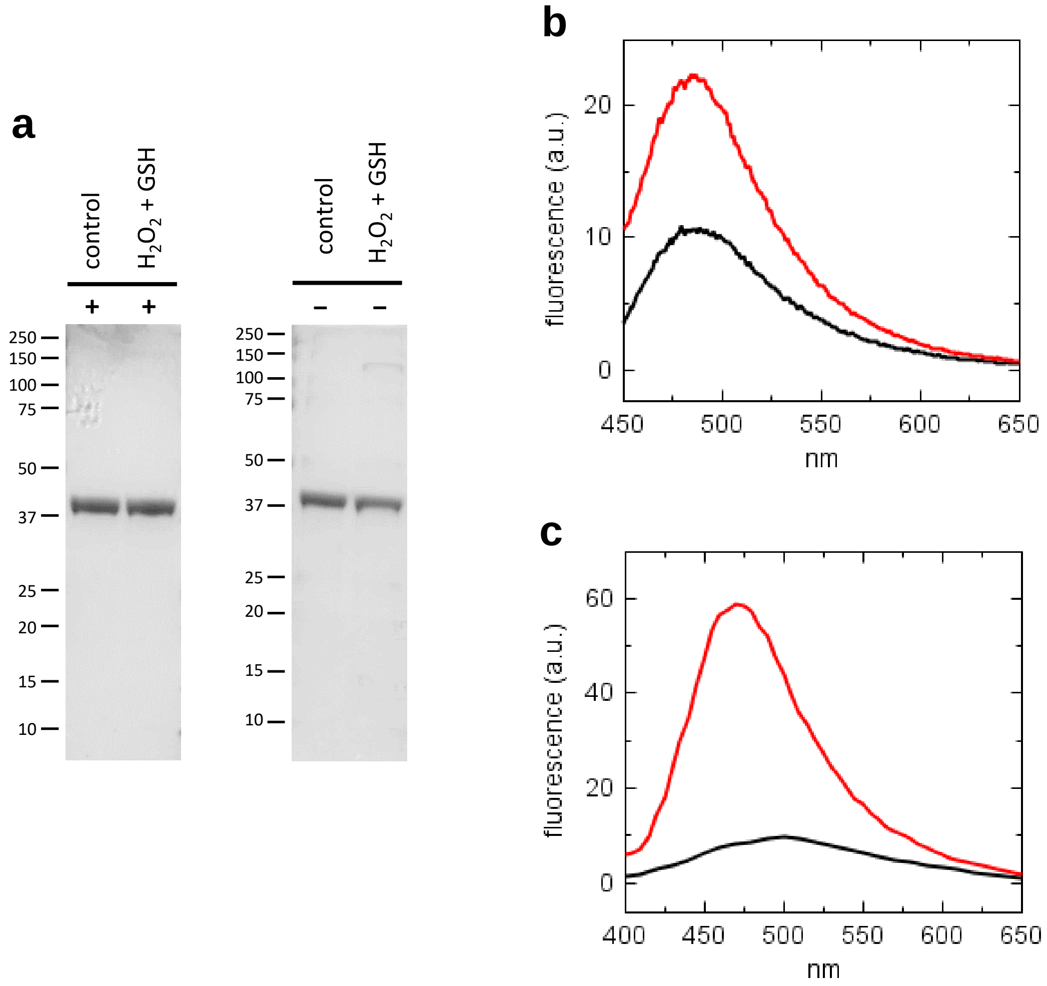

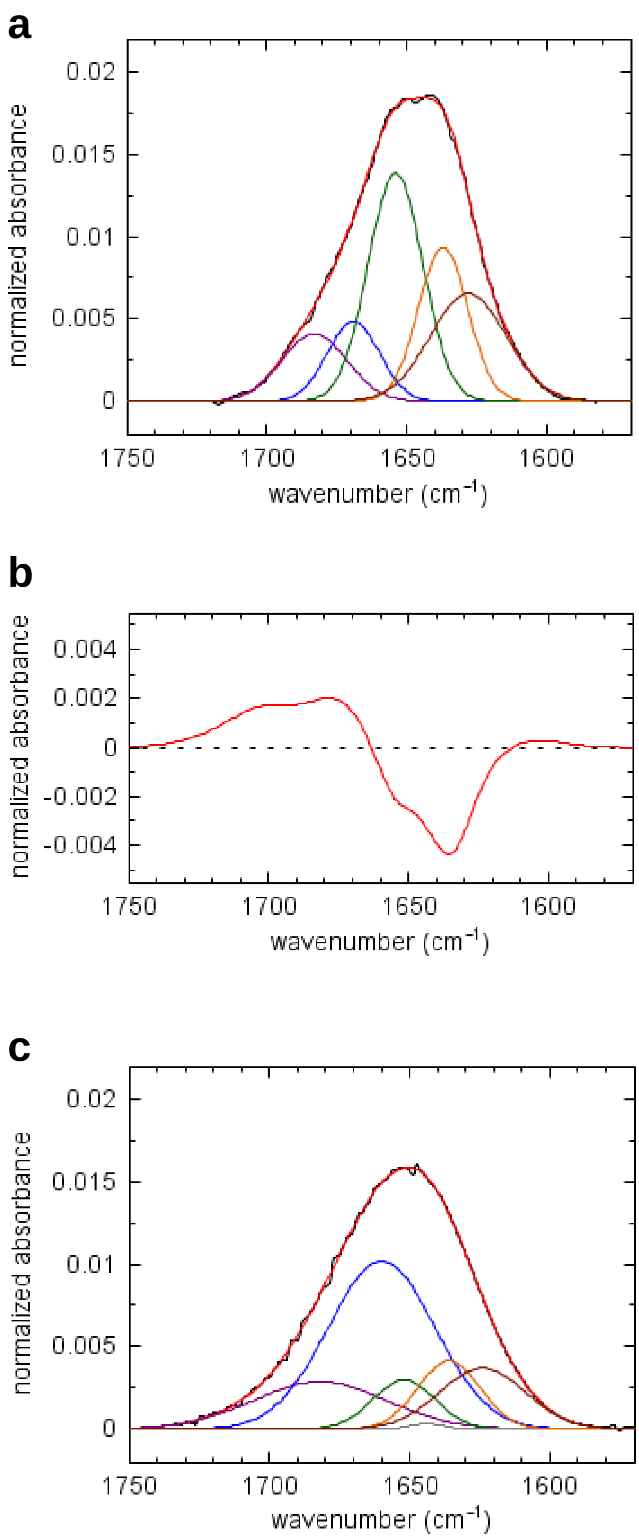

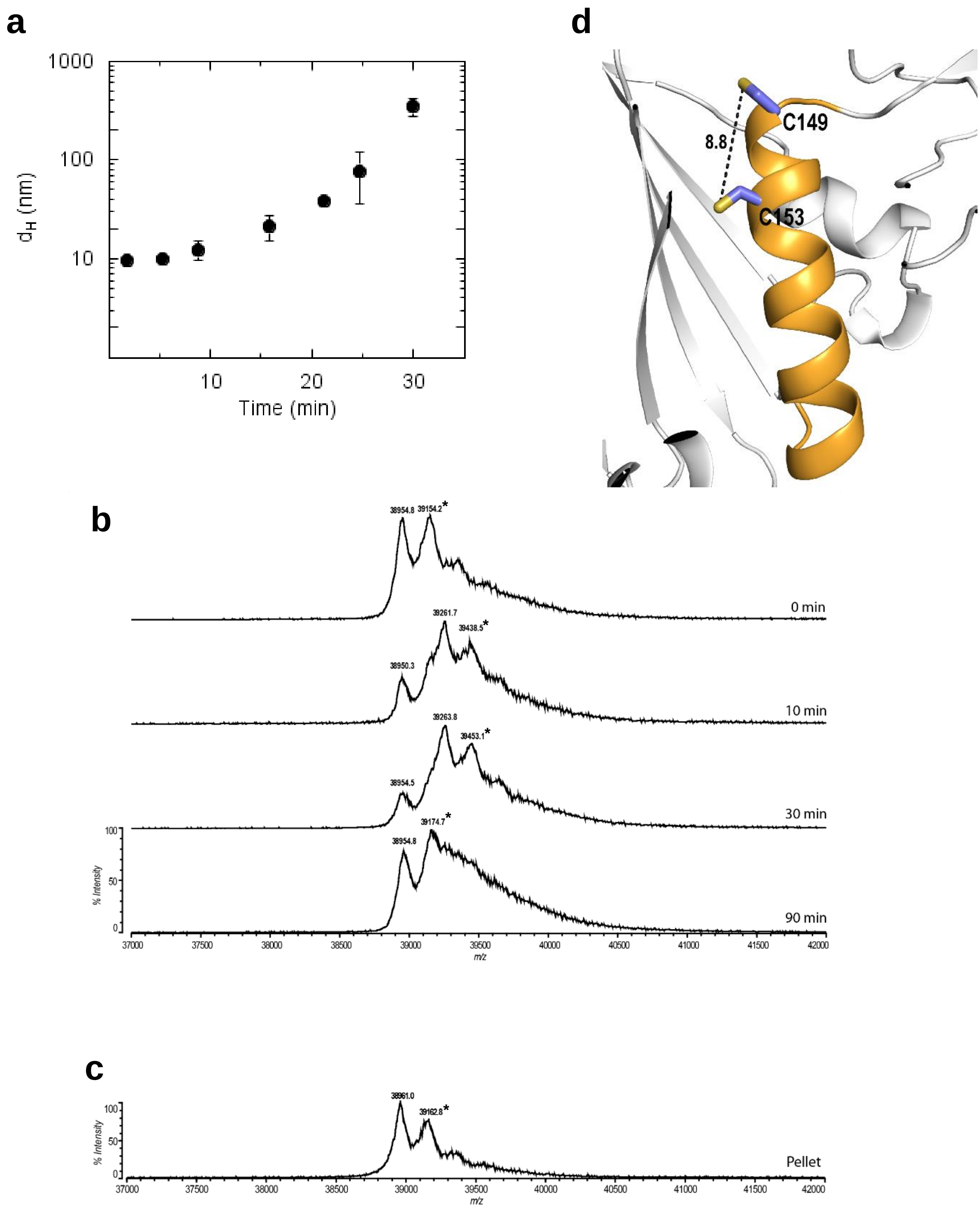

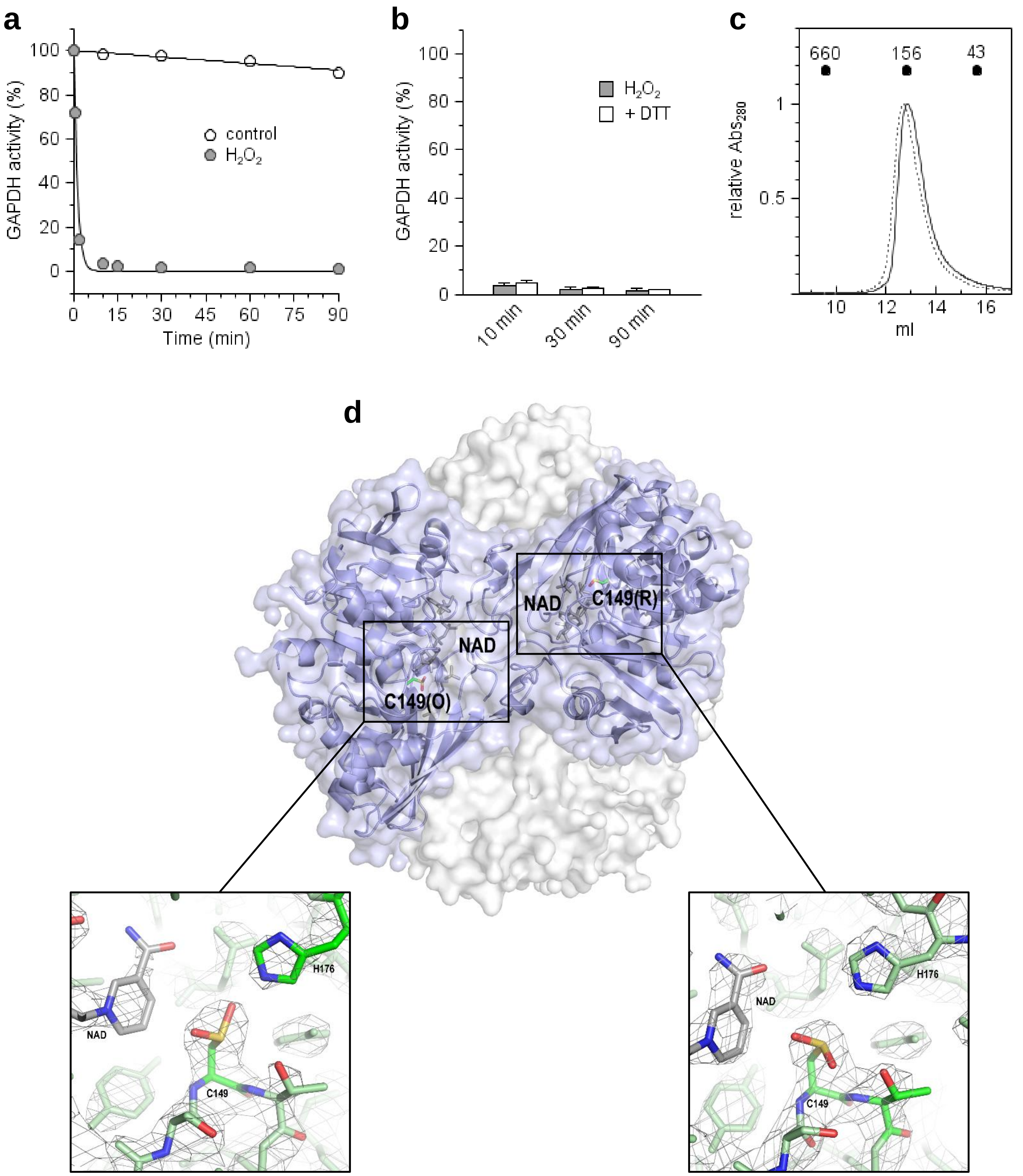

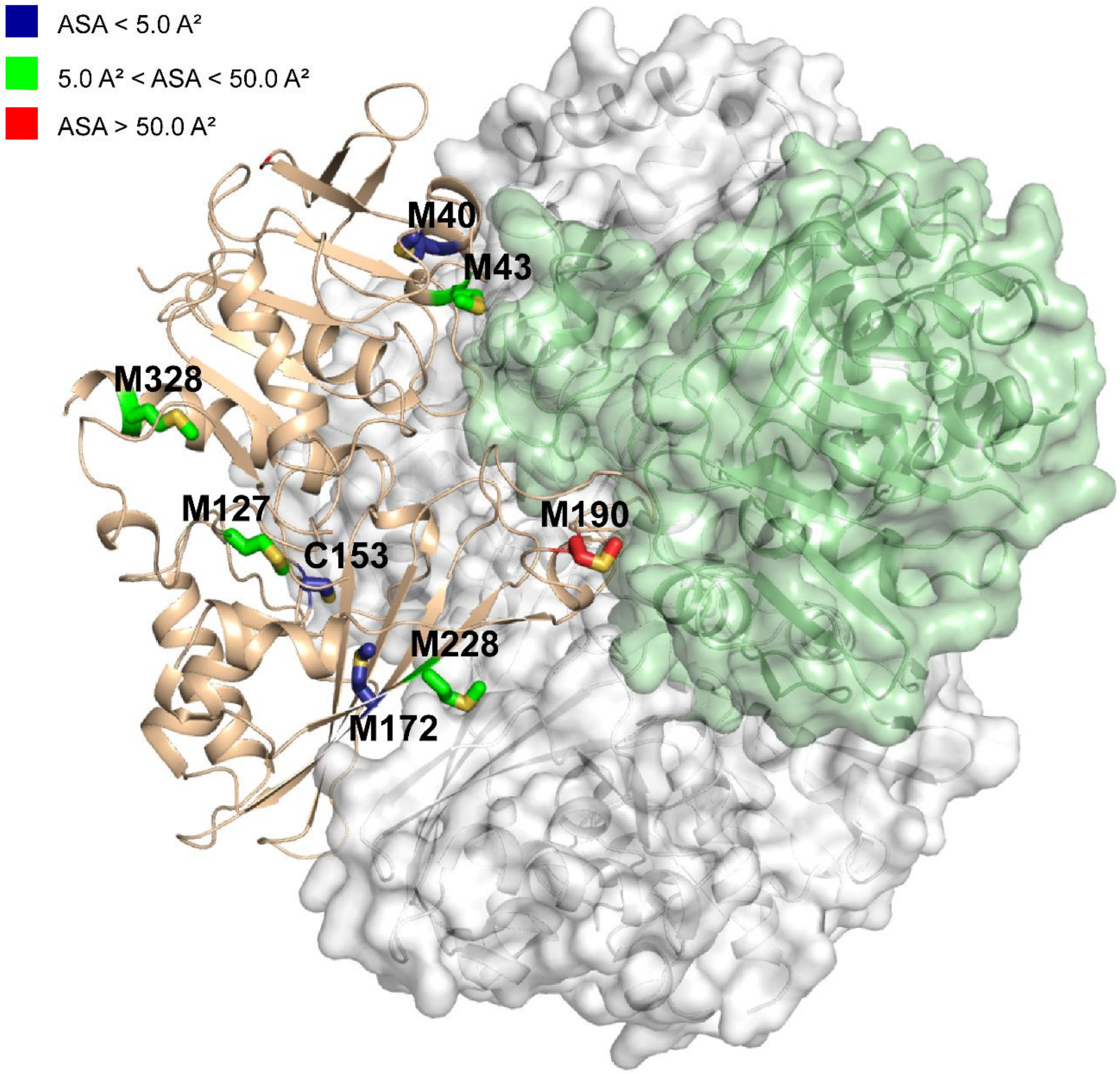

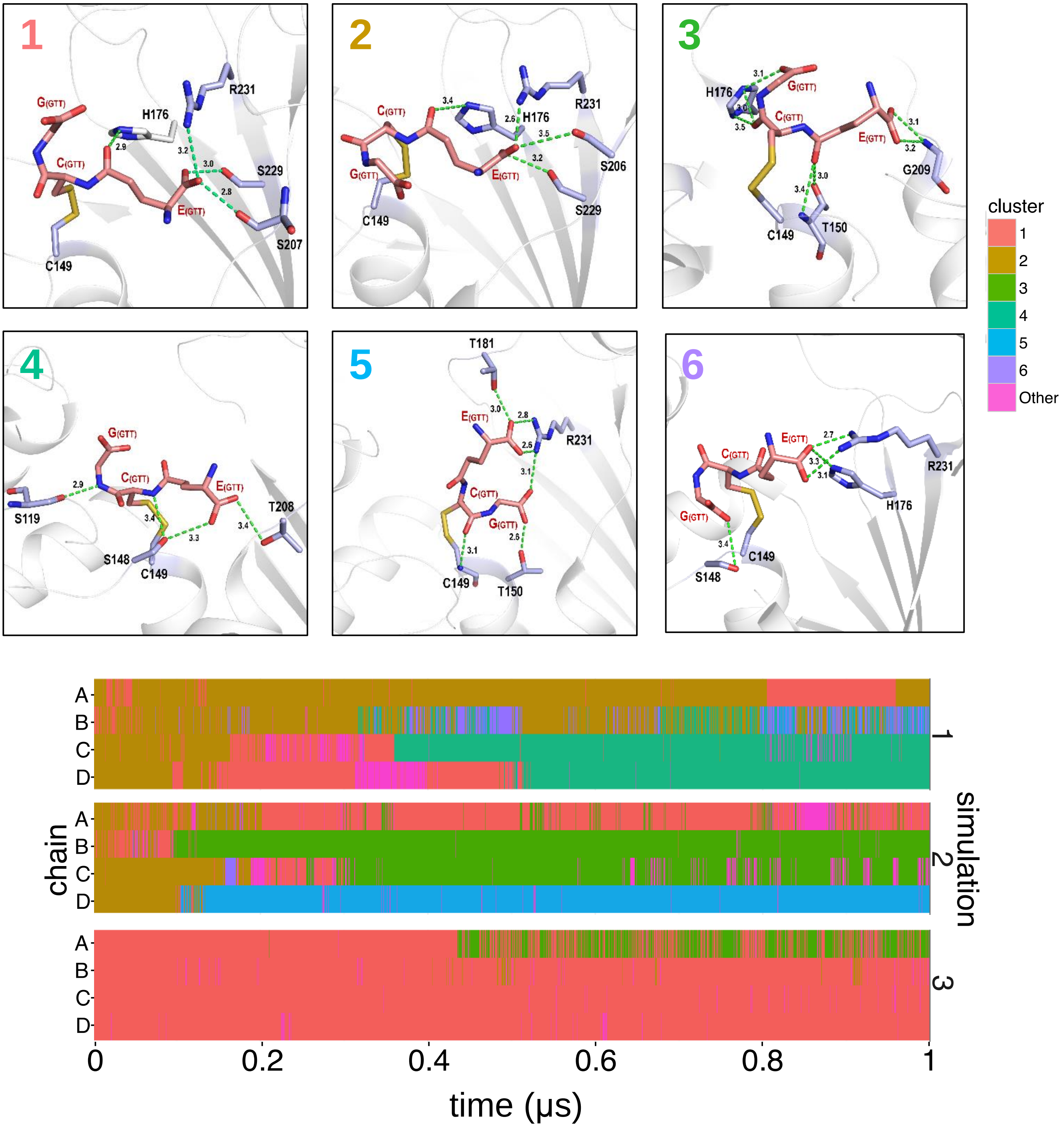

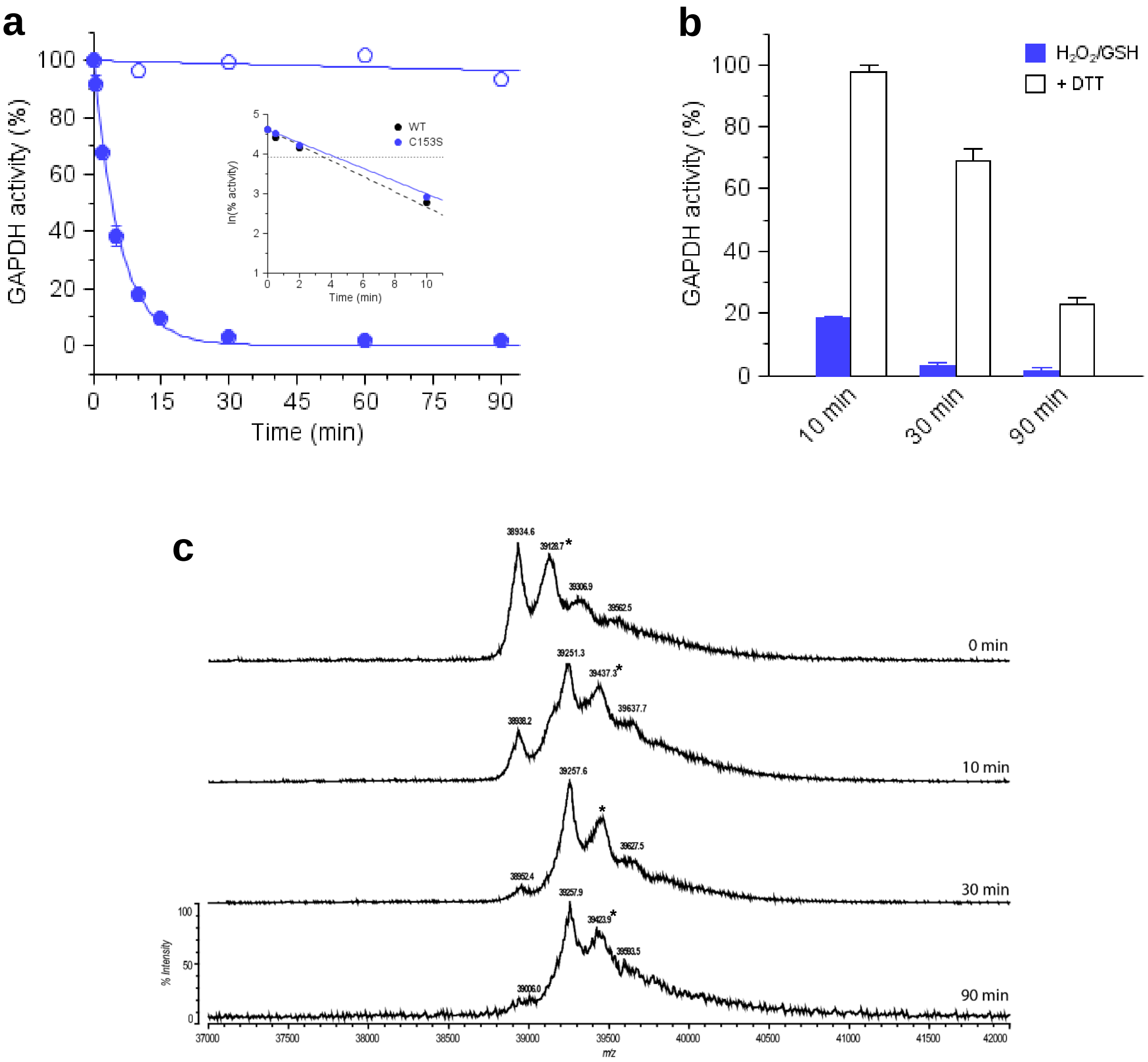
